# The brain’s “dark energy” puzzle *upgraded*: [^18^F]FDG uptake, delivery and phosphorylation, and their coupling with resting-state brain activity

**DOI:** 10.1101/2024.10.05.615717

**Authors:** Tommaso Volpi, John J. Lee, Andrei G. Vlassenko, Manu S. Goyal, Maurizio Corbetta, Alessandra Bertoldo

**Affiliations:** Department of Radiology and Biomedical Imaging, Yale University School of Medicine, New Haven, CT 06520, USA; Padova Neuroscience Center, University of Padova, 35129, Padova, Italy; Neuroimaging Laboratories at the Mallinckrodt Institute of Radiology, Washington University School of Medicine, St Louis, MO 63110, USA; Department of Neuroscience, University of Padova, 35121, Padova, Italy; Department of Information Engineering, University of Padova, 35131, Padova, Italy

**Keywords:** Brain glucose metabolism, Kinetic modeling, Microparameters, Spontaneous activity, Multilevel modeling

## Abstract

The brain’s resting-state energy consumption is expected to be mainly driven by spontaneous activity. In our previous work, we extracted a wide range of features from resting-state fMRI (rs-fMRI), and used them to predict [^18^F]FDG PET SUVR as a proxy of glucose metabolism. Here, we expanded upon our previous effort by estimating [^18^F]FDG kinetic parameters according to Sokoloff’s model, i.e., *K*_i_ (irreversible uptake rate), *K*_1_ (delivery), *k*_3_ (phosphorylation), in a large healthy control group. The parameters’ spatial distribution was described at a high spatial resolution. We showed that while *K*_1_ is the least redundant, there are relevant differences between *K*_i_ and *k*_3_ (occipital cortices, cerebellum and thalamus).

Using multilevel modeling, we investigated how much of the regional variability of [^18^F]FDG parameters could be explained by a combination of rs-fMRI variables only, or with the addition of cerebral blood flow (CBF) and metabolic rate of oxygen (CMRO_2_), estimated from ^15^O PET data. We found that combining rs-fMRI and CMRO_2_ led to satisfactory prediction of individual *K*_i_ variance (45%). Although more difficult to describe, *K*_i_ and *k*_3_ were both most sensitive to local rs-fMRI variables, while *K*_1_ was sensitive to CMRO_2_. This work represents the most comprehensive assessment to date of the complex functional and metabolic underpinnings of brain glucose consumption.

## Introduction

The complex interplay between the brain’s metabolic rate of glucose (CMRglc) and oxygen (CMRO_2_), cerebral blood flow (CBF), and brain activity has been the subject of investigation for a long time^1,2,3^. One of the most intriguing findings in neuroscience has been the key role that spontaneous activity plays in the context of neural metabolism. As most brain energy consumption in terms of glucose and oxygen happens during rest^4,5^, one would expect some degree of coupling between indices of brain metabolism (CMRglc, CMRO_2_), blood flow, and measures of resting-state brain activity, such as those which can be derived from resting-state blood oxygen level dependent (BOLD) functional MRI (rs-fMRI)^6^. A moderate-to-strong association between resting-state CMRglc, CMRO_2_ and CBF has been postulated, and demonstrated by PET studies using [^18^F]FDG, [^15^O]H_2_O, and [^15^O]O_2_ tracers^7,8^. A growing number of studies have also tested the coupling between brain glucose metabolism and features obtained from rs-fMRI, detecting complex associations, with stronger coupling of glucose metabolism with local indices of BOLD activity and synchronization, and marked between-individual variability in the strength of the association^8,9,10,11,12,13^.

In our previous work^13^, we explored the link between the spatial topography of glucose metabolism, as measured via [^18^F]FDG PET, and a variety of rs-fMRI measures in the healthy brain, attempting to capture the complexity of brain metabolism through a multifaceted assessment of spontaneous activity. As in many other research works^10,11,12^, we employed a semi-quantitative measure of [^18^F]FDG uptake, i.e., the standardized uptake value ratio (SUVR). While this was validated to be a reliable index of glucose consumption in the healthy brain, it gives a simplified view of the physiological events that can be tracked by [^18^F]FDG PET: this tracer, if combined with compartment modeling, can be used to separately estimate tracer delivery (*K*_1_) across the blood-brain barrier (BBB) mediated by glucose transporters (GLUT1, GLUT3)^14^, clearance into the venous blood (*k*_2_), and phosphorylation rate by hexokinase (*k*_3_). These microparameters (*K*_1_, *k*_2_, *k*_3_) complement and enrich the picture given by the irreversible uptake rate of [^18^F]FDG (*K*_i_ = *K*_1_*k*_3_/(*k*_2_*+k*_3_)), which is a composite macroparameter^15,16^. *K*_1_ is an expression of both CBF and the tracer’s extraction fraction *E* (*K*_1_ = *E* · CBF), and thus BBB permeability^17^. *k*_3_ represents glucose phosphorylation, which is the rate limiting step for glucose utilization, and thus of particular physiological and pathological relevance^18,19^; *k*_3_ should be strongly related to *K*_i_, making *K*_i_ a good proxy of glucose phosphorylation events, but there are regions where removing the impact of delivery may prove relevant. The spatial distribution of the microparameters was investigated for the first time in the 1980’s^20^, and in few other works^19,21^, but a more fine-grained evaluation is warranted, considering lingering questions in the fields of neural metabolism and spontaneous activity.

In this work, we focused on fully exploiting the physiological information that can be extracted from dynamic [^18^F]FDG PET data in a large dataset of healthy controls (n = 47). This was intended to expand upon our previous study, where, using two datasets, we showed that rs-fMRI measures can explain a moderate portion of regional metabolic variability (as described by SUVR), mainly driven by local rs-fMRI features^13^. However, a significant amount of variance remained unexplained. Here, we evaluated the spatial distribution of [^18^F]FDG *K*_i_, *K*_1_, and *k*_3_, and related them to a multitude of potential rs-fMRI predictors (**Table 1**), divided into four pools^13^: 1) *signal*, 2) hemodynamic response function (*HRF*)^22^, 3) static functional connectivity (*sFC*)^23^, 4) time-varying functional connectivity (*tvFC*)^24^. We also related *K*_i_, *K*_1_, *k*_3_ to CBF and CMRO_2_, as estimated from [^15^O]H_2_O, [^15^O]O_2_ PET, which more directly reflect hemodynamics and oxidative metabolism^25^. The main aims driving our analyses were:

1. assessing the spatial distribution of [^18^F]FDG *K*_i_, *K*_1_, *k*_3_ across brain regions in healthy individuals, with particular focus on *K*_1_ and *k*_3_, assuming they would provide unique information;
2. evaluating how much the individual-level spatial coupling between rs-fMRI measures (as proxies for the properties of spontaneous brain activity) and [^18^F]FDG PET changes when considering macro-and microparameters instead of SUVR, employing multivariable mixed-effects modeling (MEM), and the same functional features identified by feature selection in our SUVR model^13^, to test their reproducibility and generalizability;
3. evaluating the role of CBF and CMRO_2_ when added to the fMRI-based description of [^18^F]FDG kinetic parameters, hypothesizing that only through this more comprehensive assessment of the brain’s functional properties we would reach a somewhat satisfactory description of glucose metabolism in its entirety.

**Table 1.**
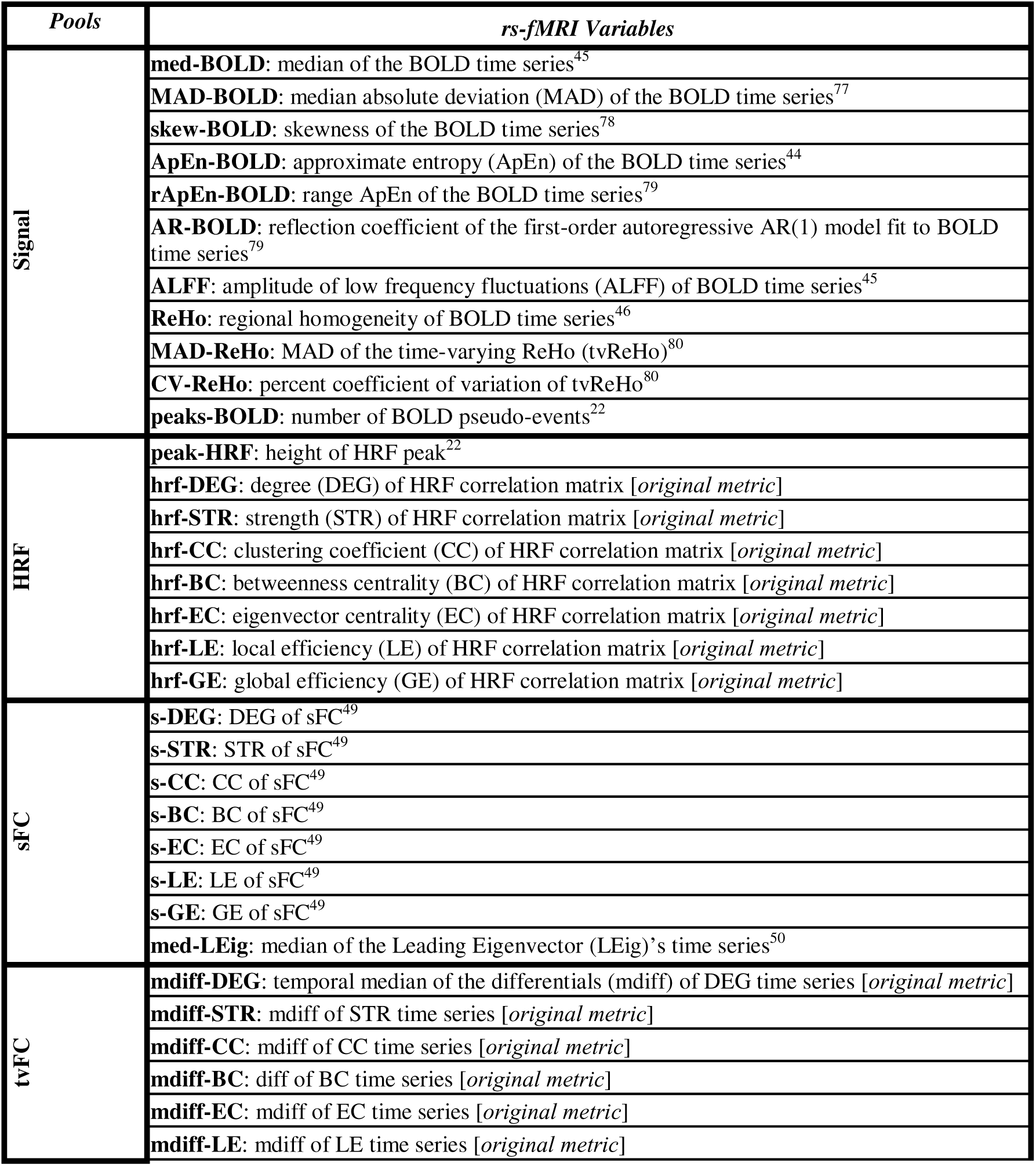
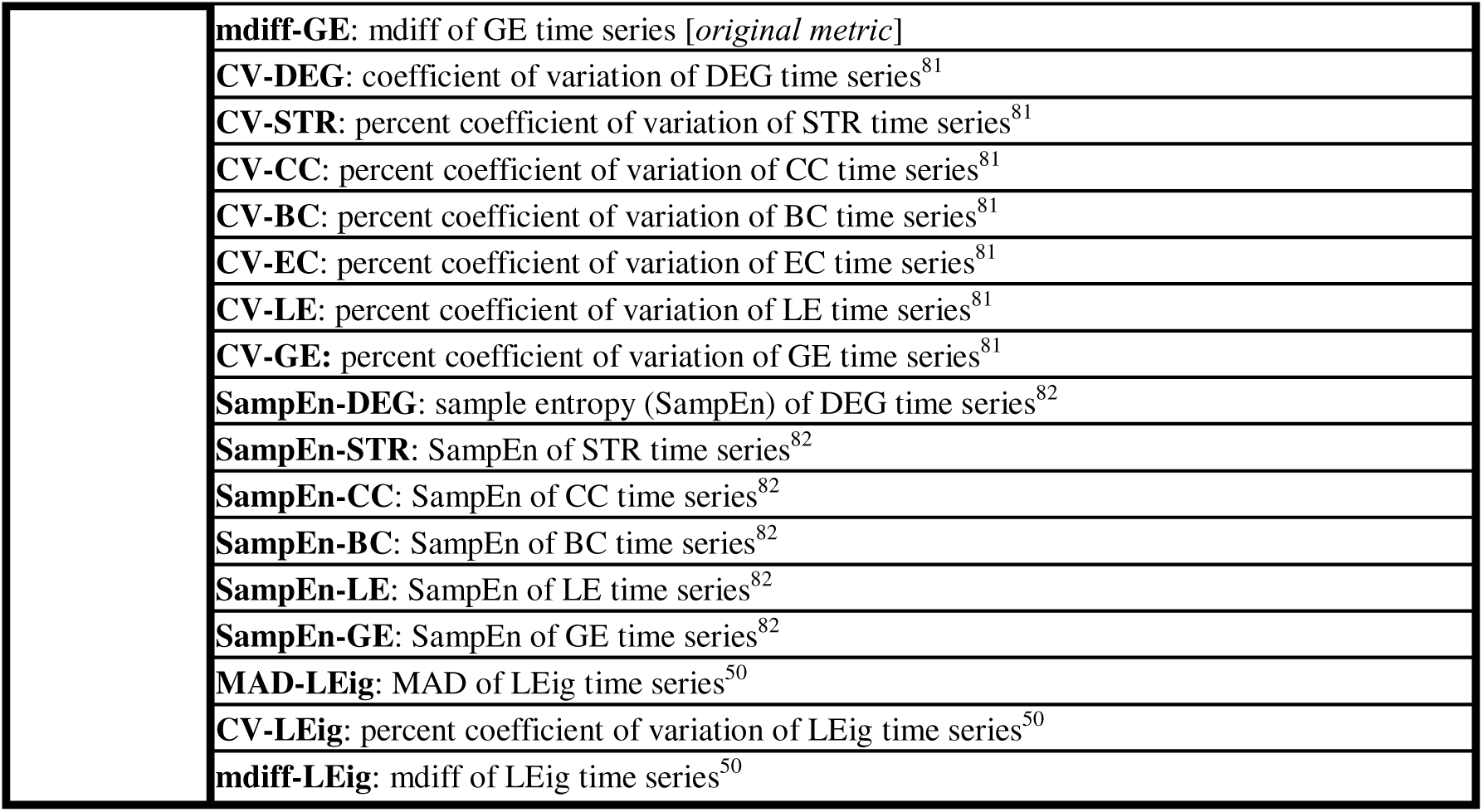
Extracted rs-fMRI features and their categories. Fifty fMRI-derived variables, divided according to the pool to which they belong: 1) signal, 2) hemodynamic response function (HRF), 3) static functional connectivity (sFC), 4) time-varying functional connectivity (tvFC). See **Supplementary Methods** for full description of the features.

The main scheme of the analysis is reported in **Figure 1**.

**Figure 1:**
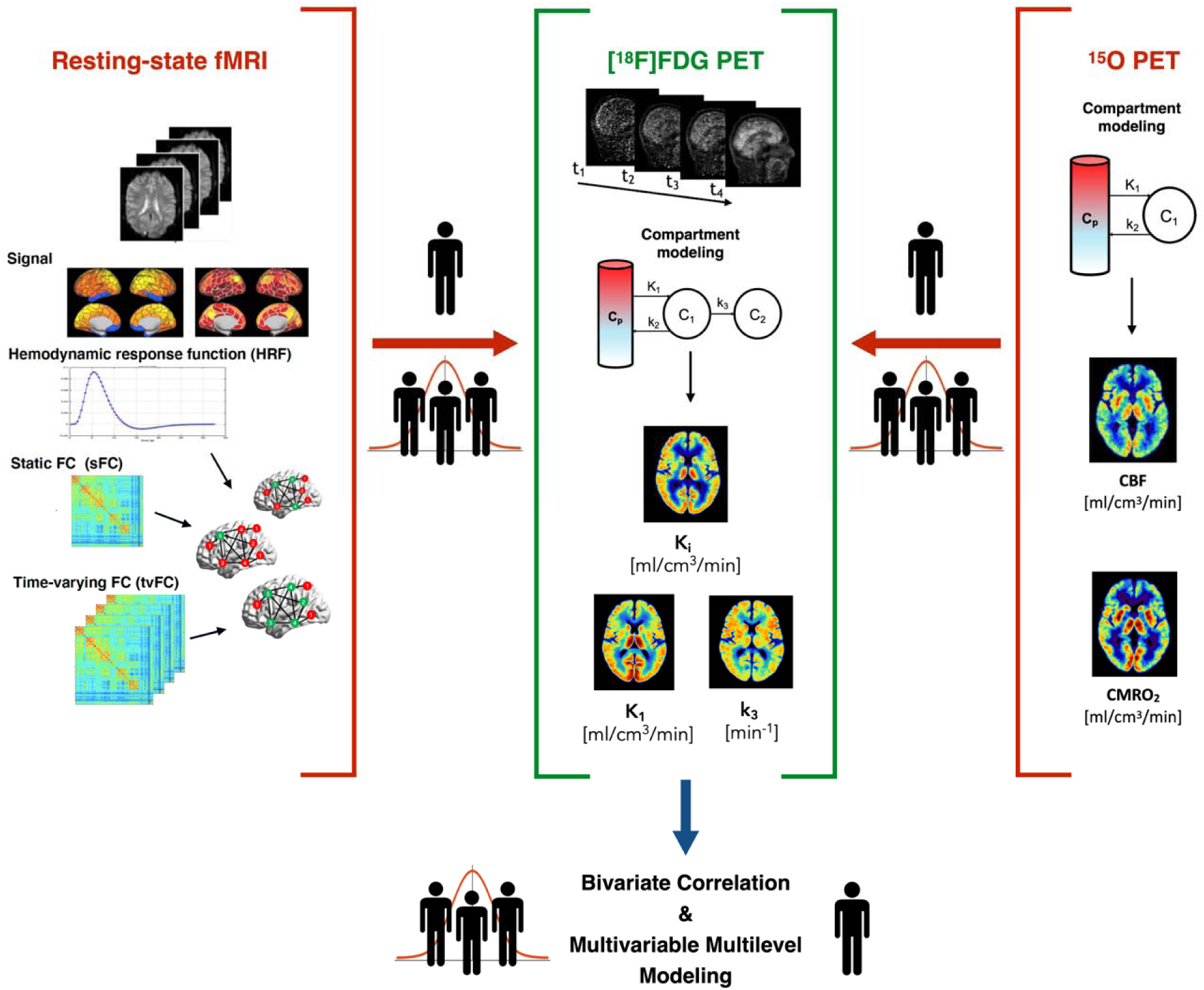
Flowchart of the analysis. The rs-fMRI data (*left*) were used to extract fifty features representative of four “pools”, i.e., 1) *signal* and *local* measures, 2) hemodynamic response function (*HRF*), 3) static functional connectivity (*sFC*), 4) time-varying functional connectivity (*tvFC*), which were parceled into 216 ROIs. [^18^F]FDG dynamic PET data (*center*) were fitted with an irreversible two-tissue compartment model to obtain voxel-wis estimates of [^18^F]FDG kinetic parameters (most importantly, for the purpose of the analyses, *K*_i_, *K*_1_ and *k*_3_), which were also parceled. [^15^O]H_2_O and [^15^O]O_2_ dynamic PET data (*top right*) were also quantified using a reversible one-tissue compartment model to obtain estimates of CBF and CMRO_2_, which were also parceled. The spatial coupling across ROIs between *predictors* (A) rs-fMRI features, B) CBF and CMRO_2_) – marked in *red*, and *outcomes* ([^18^F]FDG kinetic parameters *K*_i_, *K*_1_ and *k*_3_) – marked in *green*, was investigated on two levels, i.e., at group-averag level (*three-persons icon*) and at individual level (*one-person icon*) via bivariate correlation and multivariable multilevel modeling.

## Materials and Methods

### Participants and imaging protocols

The dataset includes 47 healthy participants (22 F; 57.4 ± 14.8 years) from the Adult Metabolism & Brain Resilience study^26^. Imaging procedures were approved by Human Research Protection Office and Radioactive Drug Research Committee at Washington University in Saint Louis. T1w and rs-fMRI (TR/TE=800/33 ms, 2.4 mm isotropic, multiband factor 6) were acquired on a Siemens Prisma^fit^ scanner. Eyes-closed [^18^F]FDG (60 min), [^15^O]H_2_O (3 min), [^15^O]O_2_ (3 min) PET acquisitions were performed on a Siemens ECAT EXACT HR+.

All studies were conducted according to the principles outlined in the Declaration of Helsinki. All participants provided written consent. Details are reported in **Supplementary Methods** and in ^26^.

### PET kinetic modeling

Dynamic [^18^F]FDG PET data were motion-corrected^27^. To perform kinetic modeling, an image-derived input function (IDIF) was extracted from dynamic PET data using a semi-automatic pipeline^28^ including 1) carotid artery segmentation; 2) selection of “hot voxels” within the mask; 3) parametric clustering^29^ to derive the raw IDIF; 4) IDIF model fitting; 5) Chen’s spillover correction^30^ using three late venous samples. Voxel-wise estimation of Sokoloff’s model parameters (irreversible two-tissue compartment model) was performed using a variational Bayesian approach^31^: k-means clustering was applied to dynamic data to extract 6 gray matter (GM) and 5 white matter (WM) clusters (based on T1w tissue segmentations), then weighted nonlinear least squares was used to estimate Sokoloff’s model parameters at the cluster level, and finally voxel-wise estimation was performed using Variational Bayesian inference based on prior distributions derived from cluster-wise estimates^32^. Parametric maps of *K*_1_ [mL/cm^3^/min] (tracer inflow), *k*_2_ [min^-1^] (efflux), *k*_3_ [min^-1^] (phosphorylation), *V*_b_ [%] (blood volume fraction) were obtained for each individual. The parametric map of *K*_i_ [mL/cm^3^/min] (irreversible tracer uptake) was obtained as *K*_i_ = *K*_1_*k*_3_/(*k*_2_*+k*_3_). The group-average maps of *K*_i_, *K*_1_, *k*_3_ are reported in **Figure 2**, while *k*_2_ and *V*_b_ are reported in **Figure S1**.

**Figure 2:**
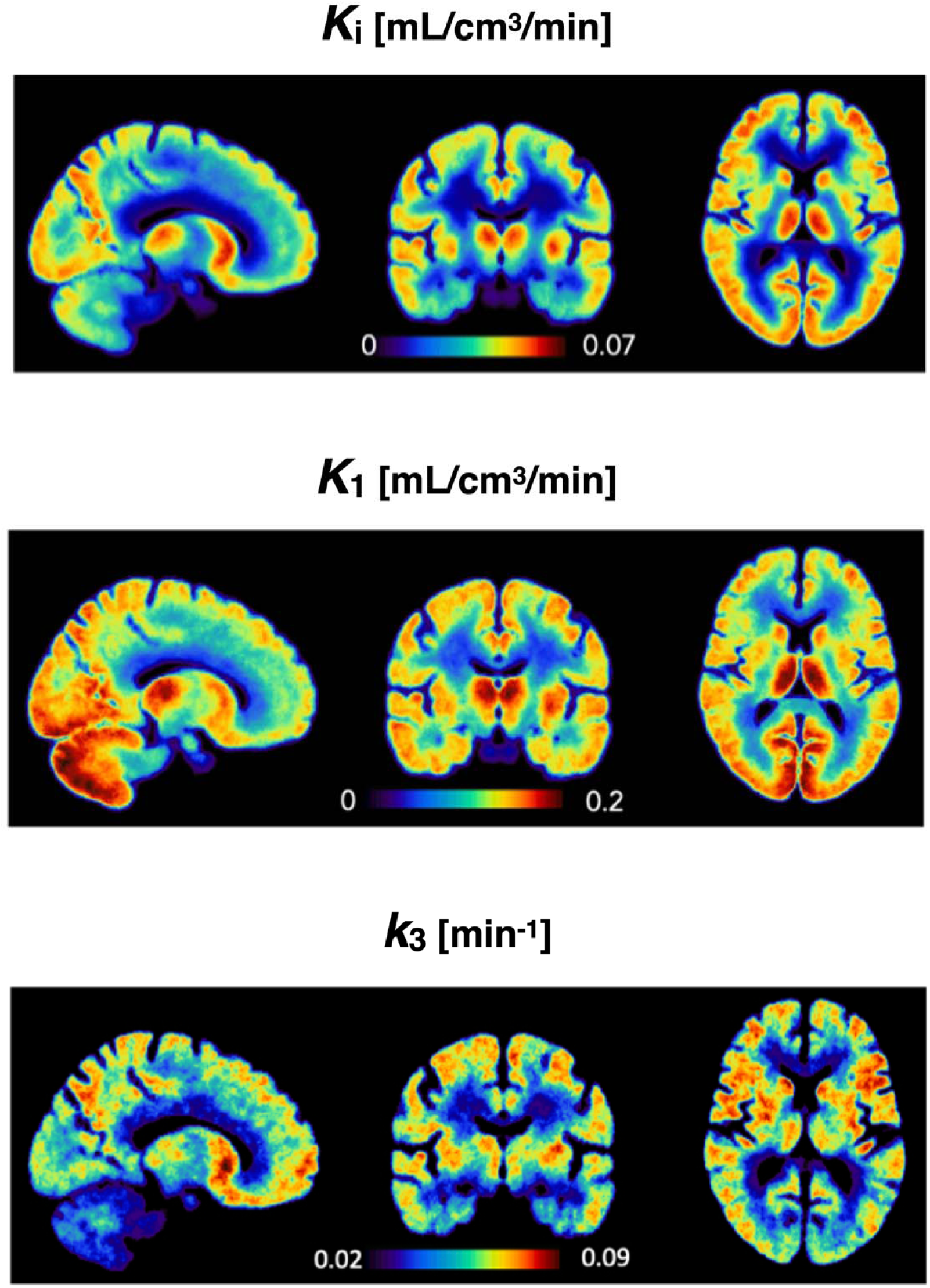
Group-average [^18^F]FDG PET parametric maps (n = 47) for *K*_i_, *K*_1_, *k*_3_ in MNI space, generated using the Variational Bayesian inference algorithm^31^.

Dynamic [^15^O]H_2_O PET was quantified with a one-tissue compartment model^33^ and a model-based input function^34^ to estimate CBF. Dynamic [^15^O]O_2_ PET was used to estimate CMRO_2_, employing a reference-tissue approach^35^. The group-average maps of CBF and CMRO_2_ are reported in **Figure S1**.

All parametric maps were parcellated into 216 regions (Schaefer atlas^36^, supplemented by 16 subcortical regions^37^), by averaging over voxels within the GM segmentation (probability > 0.8 of belonging to GM).

We did not perform partial volume correction on PET data: using the GM tissue segmentation and avoiding spatial smoothing of PET data during processing minimizes partial volume effects (PVE)^38^, as also suggested in recent work^39^. Moreover, there is no gold standard for partial volume correction, particularly in dynamic PET studies, as this procedure is known to affect kinetic modeling accuracy and potentially alter spatial metabolic patterns^40^.

Details are reported in **Supplementary Methods**.

### Resting-state fMRI processing and feature extraction

Preprocessing of rs-fMRI included slice-timing correction, nuisance regression^41,42^, high-pass filtering (cut-off = 0.008 Hz). Additional motion correction was adapted to the rs-fMRI features (i.e., despiking for tvFC, HRF, and other time-varying measures, and volume censoring for sFC and static measures^43^). The BOLD signals were also parcellated into 216 regions (GM-masked). As in ^13^, fifty rs-fMRI features were obtained in each participant. The extracted features were chosen to describe different aspects of the BOLD 1) *signal*, 2) *HRF*, 3) *sFC*, 4) *tvFC* (**Table 1**). The *signal* pool (1) relates to the basic statistics of the BOLD time series (temporal median, variance, skewness), its complexity/entropy^44^, its low-frequency fluctuations (*ALFF*)^45^, local coherence (*ReHo*)^46^ and high-amplitude events (*peaks-BOLD*). The *HRF* pool (2) relates to the HRF^22^, which links the BOLD signal to neural activity^47^, and includes the HRF peak amplitude (a potential proxy for CBF^48^), and the correlations between HRF time series of different regions, which we introduced in ^13^ to describe “purely vascular” networks, summarized at the region level by means of graph properties, for correspondence with traditional FC studies^49^. In the *sFC* pool (3) the same graph metrics are employed to characterize sFC, i.e., FC calculated across the entire fMRI scan. The *tvFC* pool (4) describes the temporal variability of graph metrics across sliding windows^24^ for each brain region. sFC and tvFC were characterized both as the correlation between BOLD signal *magnitudes*, and as the coherence of their *phases*^50^. This wide range of functional metrics were selected to represent the majority of known properties of the rs-fMRI signal and its FC. See **Supplementary Methods** for details.

### The spatial distribution of [^18^F]FDG kinetic parameters and their relationships

To investigate the spatial distribution and regional variability of *K*_i_, *K*_1_ and *k*_3_, the group-average ROI values were calculated. The *K*_i_, *K*_1_ and *k*_3_ ROI values were grouped into Schaefer’s 17 functional networks^36^. The top and bottom 20% values of each kinetic parameter were identified as ‘high’ and ‘low’, and visualized. Across-*region* Pearson’s correlations (p<0.05) between the three group-average parameter vectors were computed. Linear regression models (*K*_i_ as predictor, *K*_1_ and *k*_3_ as separate outcomes) were estimated and their weighted residuals visualized to assess the presence of regional mismatches between parameters.

### Bivariate associations between [^18^F]FDG kinetic parameters and functional features

The bivariate spatial relationship between [^18^F]FDG kinetic parameters and A) rs-fMRI properties, B) CBF and CMRO_2_, was assessed at group level, taking the region-wise mean values across individuals for each of the features. After testing for Gaussianity (p>0.05, Shapiro-Wilk test^51^), the bivariate associations across ROIs were evaluated via Pearson’s correlation (p<0.05). Differences between [^18^F]FDG parameters in their spatial correlation with A) rs-fMRI properties, B) CBF and CMRO_2_ were tested using Steiger’s z-test for dependent correlations with one variable in common (p<0.05). The average and variability (mean ± SD) of the squared values of [^18^F]FDG vs. rs-fMRI spatial correlations (R^2^) were computed, as indices of the overall strength of functional-metabolic spatial association across fMRI variables. Differences among [^18^F]FDG parameters (factor 1) and groups of rs-fMRI features (factor 2) in the strength of [^18^F]FDG-fMRI association (R^2^) were assessed using a two-way analysis of variance (ANOVA) with unbalanced design, with [^18^F]FDG parameters as first factor (3 levels: *K*_i_, *K*_1_, *k*_3_), and groups of rs-fMRI features as second factor (4 levels: signal, HRF, sFC, tvFC). Statistical differences between pairs of means were determined using Tukey-Kramer’s multiple comparison test.

The spatial heterogeneity in the [^18^F]FDG-fMRI relationship was probed by assessing correlations iteratively across clusters of regions with increasingly high or low *K*_i_, *K*_1_ or *k*_3_. The threshold levels were determined by considering linearly *increasing* percentiles (1^st^ to 85^th^) of the [^18^F]FDG parameter distribution over all regions, and linearly *decreasing K*_i_, *K*_1_ or *k*_3_ percentiles (100^th^ to 15^th^). For each threshold level (i.e., group of chosen regions), Pearson’s correlations between [^18^F]FDG parameters and all A) rs-fMRI features, B) CBF and CMRO_2_ were calculated. The type of bivariate relationship between [^18^F]FDG and rs-fMRI properties was also tested by comparing three models, i.e., 1) linear, 2) mono-exponential, 3) power law. Model selection was performed using the residual sum of squares (RSS)^53^. See ^13^ and **Supplementary Methods** for details.

### Multivariable modeling of the functional-metabolic relationship at group level

At group-average level, we used a multilinear regression approach to assess how much of the group-average [^18^F]FDG *K*_i_, *K*_1_ and *k*_3_ could be explained by a linear combination of predictors, i.e., multiple rs-fMRI variables and/or CBF, CMRO_2_. Predictors and outcome were centered and scaled by their standard deviation (SD) across brain regions, and log-transformed, as in ^13^, to account for nonlinearities (**Supplementary Results**).

Rather than performing a new feature selection for each of the kinetic parameters, we tested whether the predictors selected for SUVR in ^13^ would generalize to *K*_i_, *K*_1_ and *k*_3_: specifically, we considered both the full SUVR model (9-parameter model, 9p) and its more parsimonious version (3-parameter model, 3p)^13^. The models’ coefficients were re-estimated. The tested models were evaluated in terms of number of predictors, adjusted R-squared (R_adj_^2^), and estimate precision (i.e., percent standard error divided by estimated value, %SE). An “adapted” version of the 9p model was generated, by excluding predictors whose estimates had %SE>100%. To assess the presence of multicollinearity, we also examined the Pearson’s correlations among predictors, the condition number of the selected features, and variance inflation factors^54^.

As a second step, the addition of CBF or CMRO_2_ to the selected fMRI-based models was evaluated in terms of goodness of fit (R_adj_^2^) and %SE.

### Full hierarchical modeling of [^18^F]FDG kinetic parameters using functional predictors

As in our previous work^13^, full *multilevel* population modeling (MEM) was implemented to characterize in a single stage both fixed effects *θ* and random effects *η* contributing to the relationship between [^18^F]FDG parameters and the selected functional variables^56^. The *individual*-level model is:

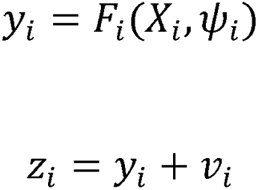

With *y_i_* as the *K*_i_, *K*_1_ or *k*_3_ prediction for the *i*^th^ individual (i = 1,,…,*m*), which is a function of *X_i_* (the fixed-effects design matrix), and of parameters *ψ_i_* to be estimated; *z_i_* is the vector of *K*_i_, *K*_1_ or *k*_3_, and *v_i_* is the *within-individual variability*, assumed to be normally distributed with zero mean and variance *σ_i_*^2^. The *population*-level model describes *ψ_i_* combining the population parameters *θ* and the random variability of individual parameters around the population mean *η*_i_:

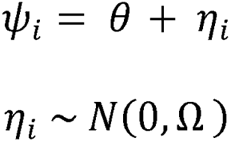

where *η*_i_ is normally distributed with zero mean, independent across individuals and with covariance matrix Ω (assumed to be full). The features selected in the previous step were included in the first-level model, i.e., either rs-fMRI features only, or including CBF or CMRO_2_, after within-individual z-scoring. The overall and individual multilevel model R^2^_adj_ were evaluated, considering the pooled data of all participants (R_adj_^2^) and the single individual data (R^2^ ), respectively. The mean (*avg. wres*) and SD of the residual unexplained variability were evaluated. The areas with higher positive ([0.5;1.5]) or negative ([-1.5;-0.5]) avg. wres were visualized to highlight where the model under-or over-estimates the outcome variable. When participant-specific covariates were available, i.e., age ([years]), sex, height ([m]), weight ([kg]), body-mass index (BMI, [kg/m^2^]), body-surface area (BSA, [m^2^]), insulin plasma levels ([mIU/L]), Pearson’s correlations (p<0.05) relating R_i_^2^ and these covariates were calculated. For more details, see **Supplementary Methods**.

## Results

### The spatial distribution of [^18^F]FDG uptake, delivery and phosphorylation rates

First, the kinetic model parameters estimated from [^18^F]FDG dynamic data, i.e., *K*_i_, *K*_1_ and *k*_3_, were evaluated (**Figure 2**). The choice to describe their spatial distribution and regional variability across the selected brain parcellation was motivated by the importance to understand the unique information they provide. The region-wise interindividual variability of [^18^F]FDG parameters is reported in **Figure S2**.

The parcels representing the top and bottom 20% values of the average regional distribution of *K*_i_, *K*_1_ and *k*_3_ were examined (**Figure 3**). Both *K*_i_ and *k*_3_ had many top nodes in lateral prefrontal areas, parietal and posteromedial cortex, while *K*_1_ had mainly a strong distribution of top posteromedial nodes including posterior cingulate and occipital areas, but also in medial sensorimotor regions. When looking at the bottom nodes, limbic areas, both at the level of the temporal poles and anterior cingulate cortex, were represented for all three parameters; however, *k*_3_ had strong presence of bottom nodes in the visual cortex, which is missing in *K*_i_, and *K*_1_ presented additional low nodes in the frontal cortex (both motor and cognitive areas) and insula. When focusing on the subcortex, we again found a similar pattern for *K*_i_ and *k*_3_, with the putamen as a top parcel, and cerebellum as a bottom one. However, the differences emerge for the caudate, which is a bottom node only for *K*_i_, and for the thalamus, which is low only for *k*_3_. In the case of *K*_1_, the putamen is again a top parcel, and the caudate is a bottom node like in the case of *K*_i_; the thalamus and cerebellum are instead amongst the top regions.

**Figure 3:**
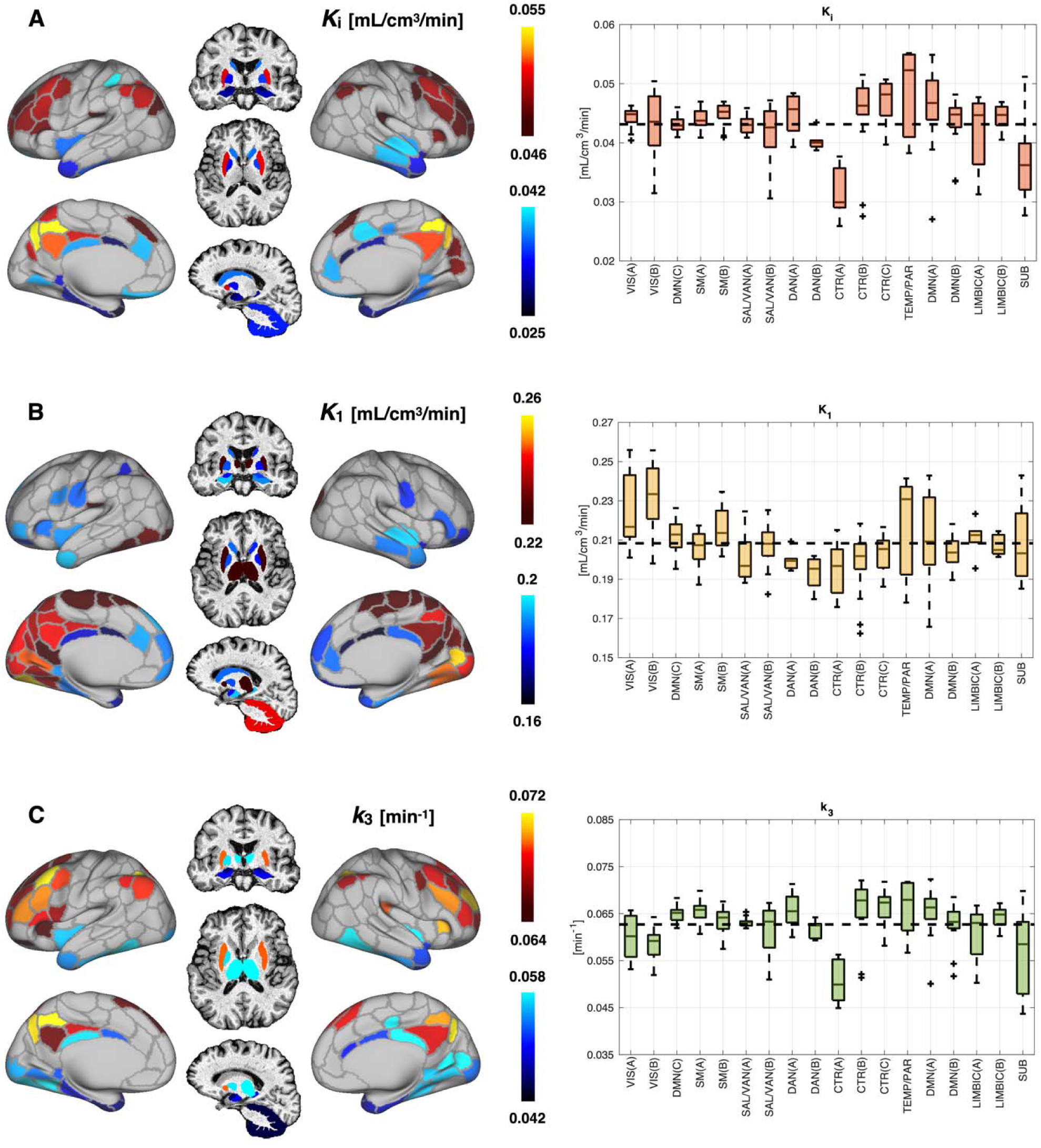
Top and bottom 20% values of group-average *K*_i_ (**A**), *K*_1_ (**B**) and *k*_3_ (**C**), along with corresponding boxplots showing group-average regional values of each parameter divided according to Schaefer’s functional brain networks.

Looking at [^18^F]FDG kinetic parameters with a perspective based on functional brain networks (**Figure 3**, boxplots) does not seem to capture a clear ranking, at least for *K*_i_ and *k*_3_, consistent with our observations for SUVR^12^. *K*_1_ is clearly highly represented in visual areas.

The spatial correlations (Pearson’s r) between the group-average parameters (**Figure S3**) across the regions of the chosen parcellation are as follows: *K*_i_ vs. *K*_1_: r = 0.56 (p < 10^-18^); *K*_i_ vs. *k*_3_: r = 0.88 (p < 10^-70^); *K*_1_ vs. *k*_3_: r = 0.19 (p = 0.006).

To better quantify the extent of the regional mismatch between *K*_i_ and each microparameter, we plotted the weighted residuals of the two linear regression models (*K*_1_ or *k*_3_ as outcome, *K*_i_ as predictor), by showing only the highest positive or negative residual values, to emphasize the regions where *K*_i_ fails to predict *K*_1_ or *k*_3_ well (**Figure S4**). *K*_1_ has higher-than-expected values in posteromedial areas, while it fails to follow the high *K*_i_ values of the lateral frontal areas and caudate nuclei. As for *k*_3_, we find markedly lower values in visual cortex and cerebellum than expected by *K*_i_, but also in thalamus; instead *k*_3_ relatively exceeds *K*_i_ mainly in the caudate nuclei, but also in insular and lateral cortical areas.

Across-individual region-wise correlations between [^18^F]FDG parameters are reported in **Figure S5**.

### Bivariate associations with functional features

As in our previous work^13^, we extracted 50 rs-fMRI variables at the individual level, and subdivided them into 4 *a priori*-defined pools: 1) *signal*, 2) *HRF*, 3) *sFC*, 4) *tvFC* (**Table 1**).

The correlation matrix between all available functional features, i.e., A) the 50 rs-fMRI features, B) CBF and CMRO_2_, at group-average level are reported in **Figure S6**.

The Pearson’s correlations between group-average [^18^F]FDG parameters and rs-fMRI features are presented in **Figure 4** (individual-level results in **Figure S7**).

**Figure 4:**
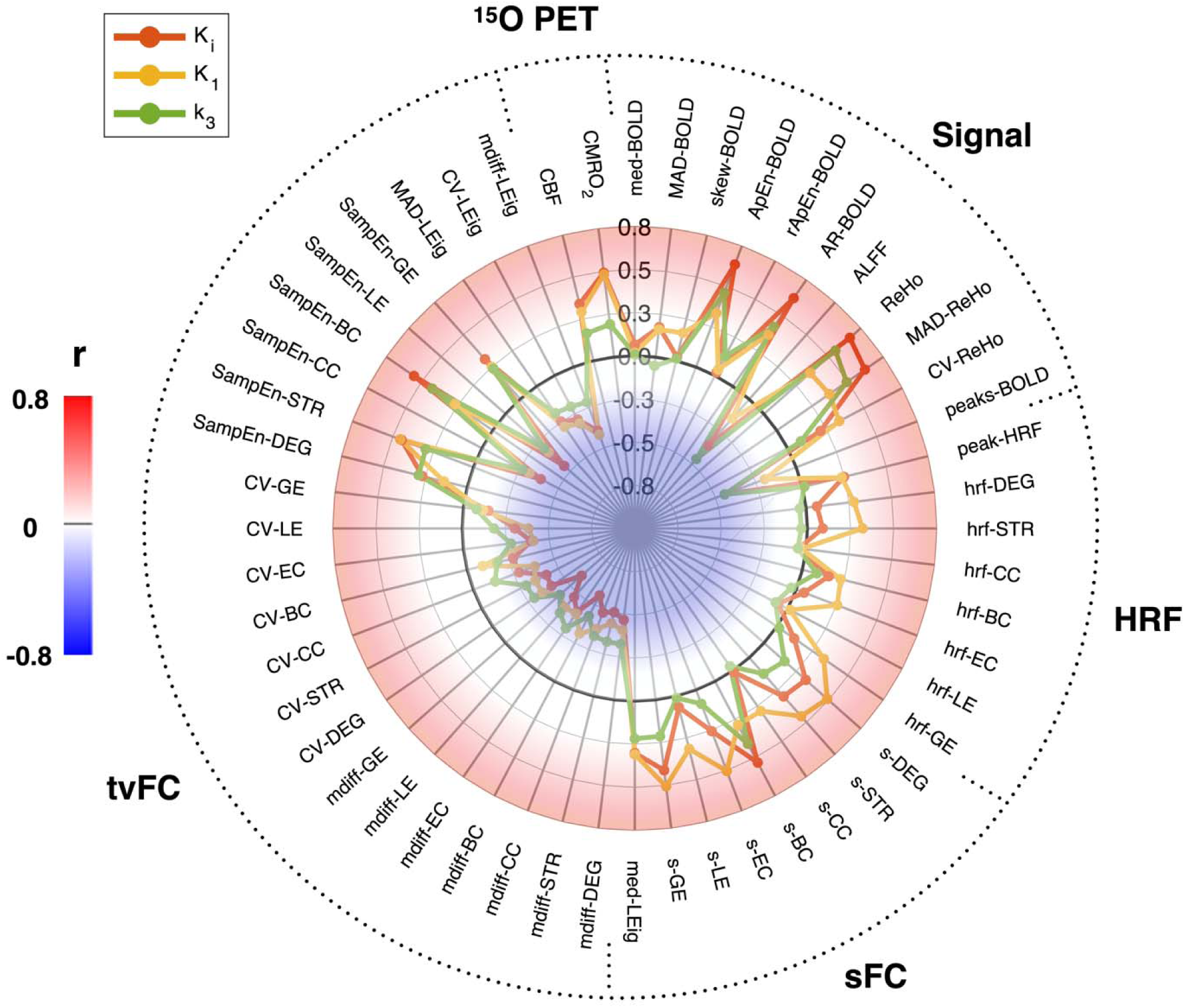
Spider plot of Pearson’s correlations across brain regions between group-average [^18^F]FDG kinetic parameters (*K*_i_, *K*_1_ and *k*_3_) vs. group-average rs-fMRI features (divided into 4 a priori defined pools: 1) signal, 2) HRF, 3) sFC, 4) tvFC), and CBF and CMRO_2_ (^15^O PET).

In the signal pool, moderate-to-strong positive or negative correlations are present for *K*_i_ and *k*_3_ with features related to rs-fMRI “signal” features (*ALFF*, *ReHo* and its temporal variability, and *peaks-BOLD*), while *K*_1_ shows weaker coupling (not significant, 2-way ANOVA). Notably, *peak-HRF*, which represents blood flow-related information, is significantly, though weakly correlated with *K*_1_ and *K*_i_, but not with *k*_3_. Moreover, HRF network features are only related to *K*_1_. Interestingly, all sFC measures display significant associations with *K*_1_, but not with *k*_3_, while *K*_i_ presents a mixed situation. Finally, in the case of the tvFC pool, the pattern of correlations is similar for the three [^18^F]FDG parameters, albeit with stronger negative correlations for *K*_i_ (not significant, 2-way ANOVA). When assessing the squared values of correlations with rs-fMRI features, *K*_i_ and *K*_1_ have similar magnitudes, while *k*_3_ has significantly lower correlations than *K*_i_ (Tukey-Kramer test, p < 0.01).

CBF is correlated with *K*_i_ (r = 0.43, p < 10^-10^), *K*_1_ (r = 0.38, p < 10^-8^), and *k*_3_ (r = 0.25, p < 10^-3^); CMRO_2_ has stronger associations than CBF with all parameters (Steiger’s test, p < 0.05), i.e., *K*_i_ (r = 0.60, p < 10^-22^), *K*_1_ (r = 0.61, p < 10^-22^), and *k*_3_ (r = 0.32, p < 10^-5^).

Building on our finding that the SUVR-fMRI coupling is stronger in lower SUVR nodes^13^, we reassessed [^18^F]FDG-fMRI Pearson’s correlations (p < 0.05) across nodes selected according to linearly increasing as well as decreasing percentiles of each [^18^F]FDG parameter’s distribution (**Figure S8**). Despite some differences between parameters, we confirmed that nonlinearities exist in the spatial relationship between [^18^F]FDG parameters and rs-fMRI features across brain regions, and that functional-metabolic associations are overall stronger in low-metabolism brain regions. Notably, sFC features have the most marked nonlinear associations with [^18^F]FDG parameters, especially *K*_i_ and *k*_3_ (**Figure S9**).

### The fMRI-only models of [^18^F]FDG kinetic parameters

We then evaluated whether the combination of rs-fMRI features we had chosen for SUVR^13^ (both 9p and 3p models) could explain the regional variability of *K*_i_, *K*_1_ and *k*_3_. The 9p model included 1) approximate entropy of BOLD (*ApEn-BOLD*), 2) range ApEn-BOLD (*rApEn-BOLD*), 3) *ReHo*, 4) coefficient of variation of ReHo (*CV-ReHo*), 5) *peaks-BOLD*, 6) local efficiency of HRF network (*hrf-LE*), 7) sFC betweenness centrality (*s-BC*), 8) median of leading eigenvectors of BOLD phase coherence (*med-LEig*), 9) coefficient of variation of betweenness centrality (*CV-BC*). Features 1-5 are from the signal pool, 6 is from the HRF pool, 7-8 from the sFC pool, and 9 from the tvFC pool (**Table 1**). The 3p model included 1) *ReHo*, 2) *CV-ReHo*, 3) *CV-BC*.

For group-average *K*_i_, the 9p model had an R^2^ of 0.68 (3p model: 0.60); such performance is very similar to group-average SUVR in our previous work (9p model: 0.69, 3p model: 0.59)^13^. For group-average *K*_1_, the 9p model had an R^2^ of 0.34 (3p model: 0.23). For group-average *k*_3_, the 9p model had an R^2^ of 0.5 (3p model: 0.44).

An adapted version of the 9p model was generated for each [^18^F]FDG parameter by removing features whose estimates had unacceptable precision (%SE>100%). This led to a 6-parameter model for *K*_i_ (*ApEn-BOLD*, *rApEn-BOLD*, *ReHo*, *CV-ReHo*, *med-LEig*, *CV-BC*; R^2^_adj_ = 0.69), an 8-parameter model for *K*_1_ (*rApEn-BOLD*, *ReHo*, *CV-ReHo*, *peaks-BOLD*, *hrf-LE*, *s-BC*, *med-LEig*, *CV-BC*; R^2^_adj_ = 0.34), a 5-parameter model for *k*_3_ (*rApEn-BOLD*, *ReHo*, *peaks-BOLD*, *s-BC*, *med-LEig*; R^2^_adj_ = 0.5).

The full MEM approach (**Figure 5**) with the features selected in the previous step (adapted 9p model) explained a significant proportion of individual-level variability in the spatial distribution of *K*_i_ (R^2^_pooled_ = 0.35), but less so in the case of *K*_1_ (R^2^_pooled_ = 0.14) and *k*_3_ (R^2^ = 0.21). The predictor *peaks-BOLD* was removed from the *K*_1_ model as its fixed effect estimate was not significantly different from zero. *ReHo* was confirmed as the most important explanatory parameter for *K*_i_ and *k*_3_ in particular (**Figure 5A**).

**Figure 5.**
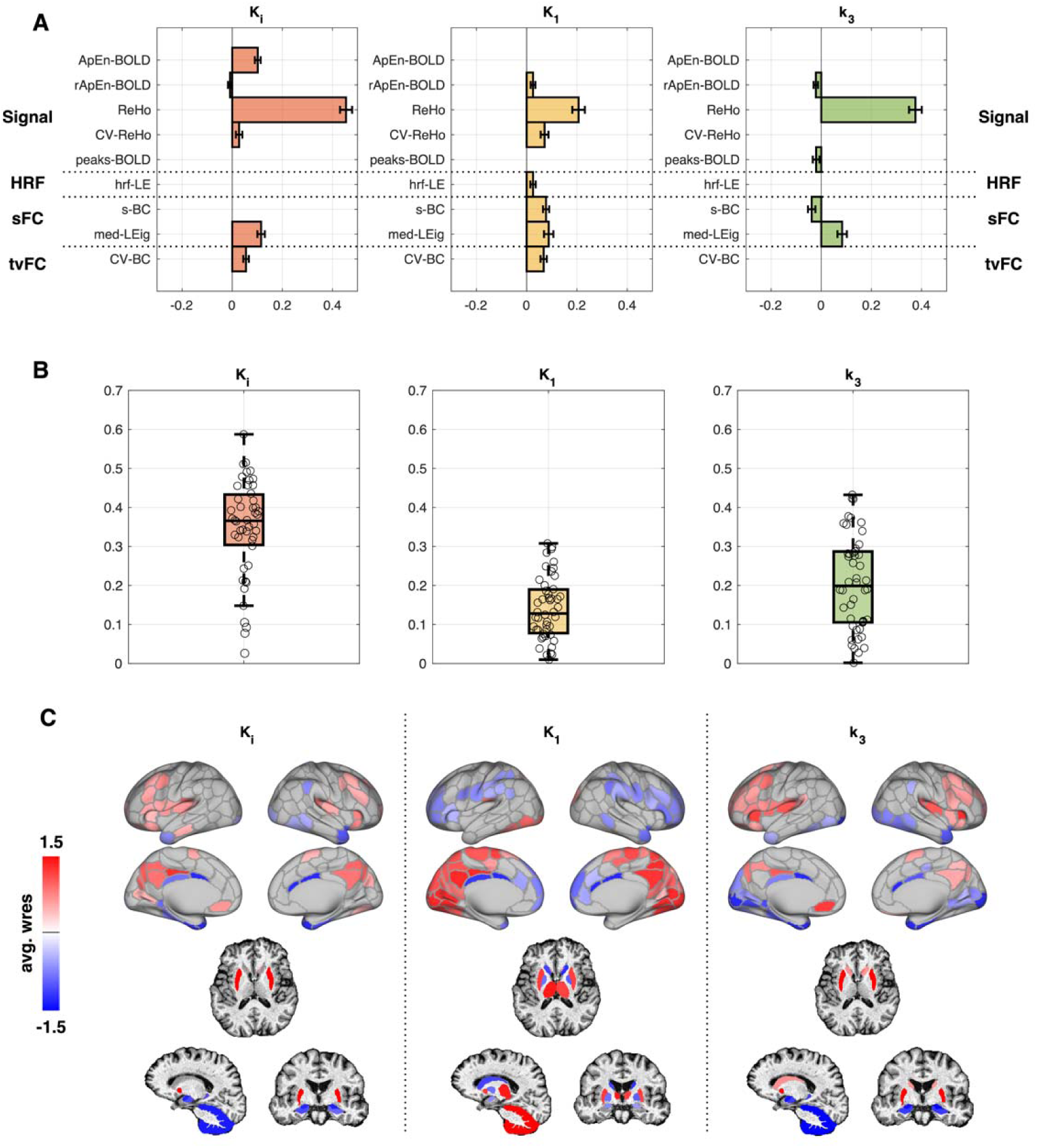
Multilevel modeling of glucose metabolism: *rs-fMRI-only models*. Fixed effects (adapted 9-parameter model) for [^18^F]FDG kinetic parameters (*K*_i_, *K*_1_ and *k*_3_): estimate weights and standard errors (SEs), which represent the parameters that best explain *K*_i_, *K*_1_ and *k*_3_ across brain regions at group level (**A**). The empty spaces correspon to features whose estimates have unacceptable precision (%SE > 100%) at group level, or estimates not significantly different from zero. Boxplots of individual R^2^ values R^2^_i_ (median and boxes of 25^th^ and 75^th^ percentile in overlay) representing the spatial variance of *K*_i_, *K*_1_ and *k*_3_ explained by the BOLD-based predictors at individual level (**B**). Across-individual average of weighted residuals of the multilevel model (*avg. wres*), visualized in the [[-1.5,-0.5];[0.5,1.5]] range for each brain region (**C**).

At the group-average level, *ReHo* explains a large proportion of variance for both *K*_i_ (R^2^ = 0.552) and *k*_3_ (R^2^ = 0.407). When recomputing the MEM estimates using only *ReHo* as a predictor, we obtained a R^2^ of 0.30 for *K*_i_ and 0.16 for *k*_3_ (0.08 for *K*_1_ ): this demonstrated that *ReHo* was responsible for the majority of the explained variance in the multivariable *K*_i_ and *k*_3_ models. Of note, in all three cases the most important parameter (with the highest associated weight) comes from the pool of signal and local rs-fMRI features.

When examining the *avg. wres* (**Figure 5C**), a marked resemblance to the top and bottom regions of each parameter (see **Figure 2**) was still present, implying that the high and low “outlier” [^18^F]FDG nodes were not adequately interpreted by the chosen rs-fMRI features. This is true especially for *K*_1_, which shows high residual values in posteromedial cortex and cerebellum, but also for *K*_i_ and *k*_3_, with high values in the putamen and low in the cerebellum.

Our finding of high between-individual variability in multilevel model R^2^ values for SUVR^13^ was confirmed here for *K*_i_, *K*_1_ and *k*_3_ (**Figure 5B**). The individual R^2^ of *K*_i_ and *k*_3_ do not correlate significantly with participants’ age or any of the peripheral metabolic information (p > 0.05). However, the R^2^ for *K* show evidence of sex difference, i.e., higher *K* R_i_^2^ for women (t-test, p < 0.01). Moreover, a negative relationship emerged between R_i_^2^ of *K* and BSA (r = -0.30, p = 0.04) and insulin levels (r = -0.35, p = 0.03) (**Figure S10**).

### Adding CBF, CMRO_2_ to models of [^18^F]FDG parameters

We then verified the impact of including CBF or CMRO_2_ along with rs-fMRI features into this multivariable modeling framework to describe the spatial distribution of [^18^F]FDG parameters. The spatial correlation between group-average CBF and CMRO_2_ is r = 0.9 (p < 10^-77^), which is why we avoided including both parameters in the same model.

At group-average level, the addition of CBF to the adapted 9p model increased the R_adj_^2^ from 0.68 to 0.78 for *K*_i_, the R_adj_^2^ from 0.34 to 0.41 for *K*_1_ , and the R_adj_^2^ from 0.5 to 0.55 for *k*_3_. The inclusion of CMRO_2_, on the other hand, increases the R_adj_^2^ from 0.68 to 0.77 for *K*_i_ , the R_adj_^2^ from 0.34 to 0.52 for *K*_1_, and the R_adj_^2^ from 0.5 to 0.51 for *k*_3_. Parameter precision remains within an acceptable range (%SE < 100%). Overall, CBF and CMRO_2_ led to similar improvements in the *K*_i_ and *k*_3_ models (moderate and minor, respectively), but CMRO_2_ importantly improved the *K*_1_ model (almost 20% of the variance).

We then assessed how these improvements impact the full MEM framework. Notably, the addition of CMRO_2_ to the adapted 9p models led to a marked increase in explained variance of the individual-level data for *K*_i_ (R^2^_pooled_: from 0.35 to 0.46) and *K*_1_ (R^2^_pooled_: from 0.14 to 0.28), with minor improvement for *k*_3_ (R^2^_pooled_: from 0.21 to 0.24). For comparison, if only CMRO_2_ is used as a predictor, we obtained a R^2^_pooled_ of 0.22 for *K*_i_ , and 0.18 for *K*_1_. The individual model R_i_^2^ can be visualized in **Figure 6B**, and the improvements appreciated by comparison with **Figure 5B**. When assessing the fixed effects, *ReHo* and CMRO_2_ had the highest weights in the *K*_i_ model, while in the *k*_3_ model, *ReHo* was the most relevant parameter, as was CMRO_2_ in the *K*_1_ model (**Figure 6A**). When examining the *avg. wres.* (**Figure 6C**), we can see the improvement in explanatory power with respect to the fMRI-only models (**Figure 5C**). This is true both for *K*_i_, which no longer shows high residual values in posteromedial cortex, and for *K*_1_, with improvements in posterior DMN, thalamus and putamen.

**Figure 6.**
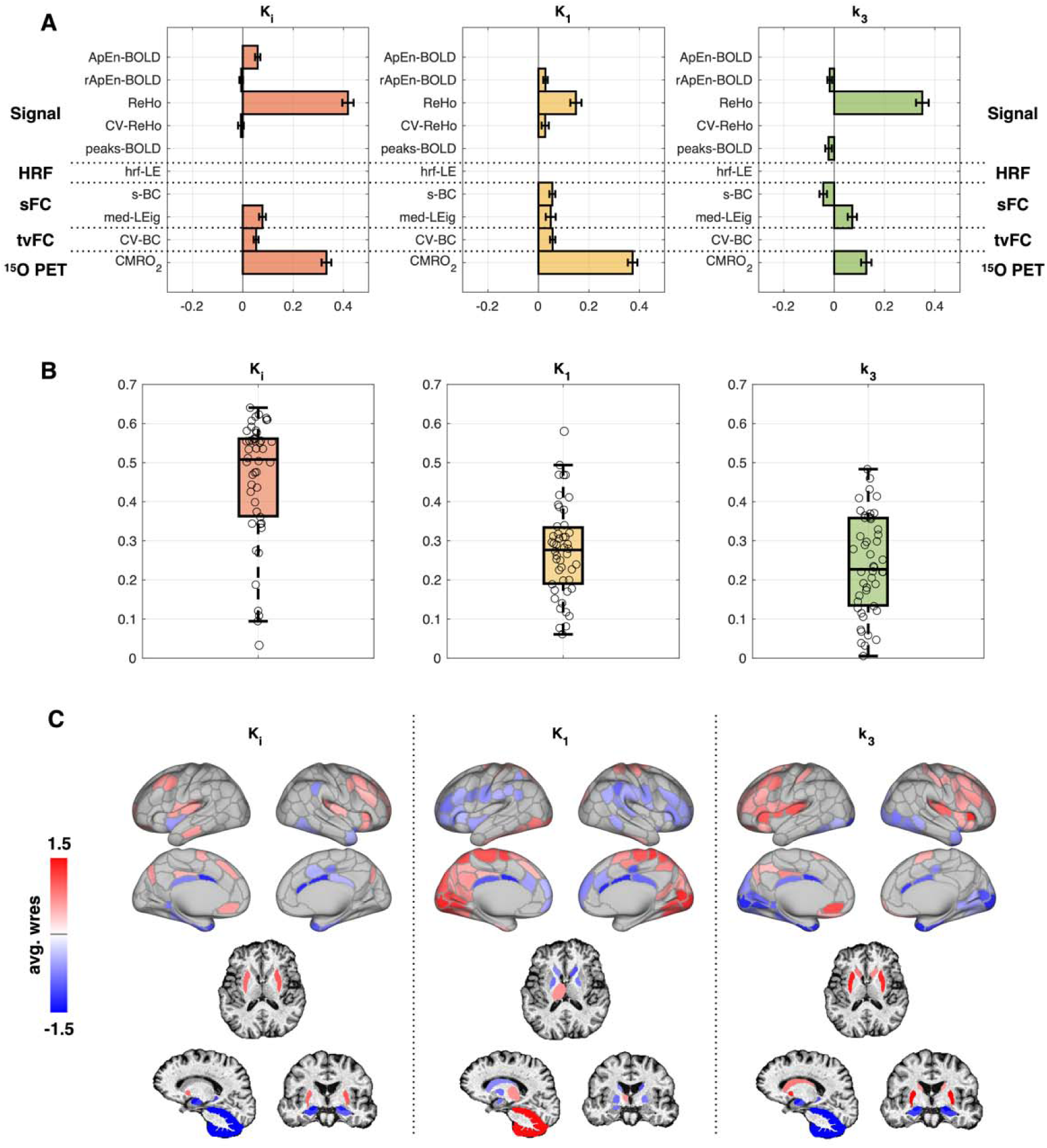
Multilevel modeling of glucose metabolism: *rs-fMRI plus CMRO_2_ models*. Fixed effects (adapted 9-parameter model + CMRO_2_) for [^18^F]FDG kinetic parameters (*K*_i_, *K*_1_ and *k*_3_): estimate weights and standard errors (SEs), which represent the parameters that best explain *K*_i_, *K*_1_ and *k*_3_ across brain regions at group level (**A**). The empty spaces correspond to features whose estimates have unacceptable precision (%SE > 100%) at group level, or estimates not significantly different from zero. Boxplot of individual R^2^ values R^2^_i_ (median and boxes of 25^th^ an 75^th^ percentile in overlay) representing the spatial variance of *K*_i_, *K*_1_ and *k*_3_ explained by the predictors at individual level (**B**). Across-individual average of weighted residuals of the multilevel model (*avg. wres*), visualized in the [[-1.5,-0.5];[0.5,1.5]] range for each brain region (**C**).

Adding CBF led to an increase in explained variance of the individual-level data for *K*_i_ (R^2^_pooled_ = 0.45), similar to CMRO_2_, while the benefit was lower for *K*_1_ (R^2^_pooled_ = 0.23) and *k*_3_ (R^2^_pooled_ = 0.25), as anticipated by the group-average modeling results.

Of note, when adding CMRO_2_ and CBF, a negative relationship between R_i_^2^ of *K* and insulin levels was still present (r = -0.41, p = 0.01).

When minimizing the number of predictor variables, i.e., only *ReHo* and CMRO_2_, we reached a R^2^_pooled_ of 0.43 for *K*_i_, and 0.21 for *k*_3_; using only *ReHo* and CBF, the R^2^_pooled_ of *K*_i_ is 0.42, and the R^2^ of *k*_3_ is 0.20.

## Discussion

In this work, we assessed the regional distribution of [^18^F]FDG kinetic parameters (*K*_i_, *K*_1_ and *k*_3_), disentangling the early steps of brain glucose metabolism (delivery vs. phosphorylation by hexokinase) for the first time at high spatial resolution. We then investigated how well these kinetic parameters can be described by a combination of features derived from rs-fMRI and expected to represent spontaneous brain activity. As in our previous work^13^, we hypothesized that a combination of many facets of spontaneous BOLD activity would be able to collectively explain (part of) glucose metabolic variance. Finally, to overcome potential limitations in rs-fMRI features’ explanatory power, we included more direct measures of hemodynamics and metabolism, i.e., regional blood flow (CBF) and oxygen consumption (CMRO_2_). We again took advantage of the multilevel modeling framework to fully account for between-individual variability in the investigated relationships. This study thus expands upon our effort^13^ to address the complexity of brain glucose metabolism^57^, which involves both oxidative and glycolytic components supporting numerous cellular processes (protein synthesis, protein modification, cell signaling, housekeeping duties, postsynaptic potentials, vesicle recycling etc.)^57,58^, with the exact partitioning of the brain’s energy budget into these processes still an object of ongoing research^4^. To better comprehend these complex biological mechanism, we used kinetic modeling to estimate [^18^F]FDG parameters at high spatial resolution (i.e., voxels, then grouped into 216 ROIs), in particular the microparameters (*K*_1_ and *k*_3_). While SUVR (which can be obtained from a short static scan, with no arterial sampling) can be a good proxy measure of *K*_i_, it is both relative and semi-quantitative, and susceptible to multiple technical and physiological biases^59,60^. We have shown here that *K*_i_, i.e., the total [^18^F]FDG uptake rate constant, is a good proxy for *k*_3_, i.e., the rate of phosphorylation events, with group-average *K*_i_ and *k*_3_ maps being highly spatially correlated. This high level of matching is expected, since [^18^F]FDG does not display flow limitation (*k*_3_ is, on average, low, and smaller than *k*_2_)^61^. However, operating at a higher level of detail than before, we have also found that *k*_3_ relatively “underestimates” *K*_i_ in visual cortex, cerebellum, and thalamus, and “overestimates” *K*_i_ in the caudate, insular and frontoparietal cortex (as shown by residuals). In such areas, especially those where *k*_3_ “underestimates” *K*_i_, tracer delivery (influx *K*_1_, and, less importantly, efflux *k*_2_) seems to play a relevant role, making *K*_i_ a biased predictor of the glucose phosphorylation events. This stresses how quantifying microparameters is not redundant, albeit more technically challenging.

When we moved to assess the explanatory power of rs-fMRI features for the parameters related to glucose utilization, i.e., *K*_i_ and *k*_3_, we found overall similar results to our findings based on SUVR^13^: a) variable degrees of spatial association with rs-fMRI variables were present, with strongest match for signal-related, local features; b) evidence of nonlinearity in this association (especially for sFC features); c) the top and bottom regions for *K*_i_ and *k*_3_ difficult to describe using only rs-fMRI features; d) marked between-individual variability in the association strength. Among these findings, the most relevant is the fact that *ReHo*, i.e., the local synchronization of the BOLD signal, emerged again as the rs-fMRI variable having the strongest spatial match with [^18^F]FDG *K*_i_ and *k*_3_. *ReHo* alone was capable of explaining 55% of *K*_i_ variance at group level, and 30% on individual data (as for SUVR); for *k*_3_, it explained 40% of group-level variance (18% at individual level). This confirms the moderate-to-strong spatial coupling between glucose metabolism, here assessed through full kinetic modeling, and measures of *local* BOLD signal coherence. It remains to be seen whether this mostly reflects the metabolic demand of brain *structure*, or rather of spontaneous *activity* levels^13^.

The next step was to include more direct hemodynamic (CBF) and metabolic (CMRO_2_) measures into this equation, which allow to describe blood flow and oxidative vs. non-oxidative brain metabolism. Notably, the strength of spatial association between these parameters and [^18^F]FDG *K*_i_ or *k*_3_ was somewhat weaker than expected, especially for CBF^8,62^. However, recent literature reports a moderate (r = 0.56), nonlinear association between fully quantitative CMRglc and CBF estimates (higher-than-expected CBF in thalamus, cerebellum and medial temporal lobe^39^), which is in line with our findings. Methodological differences in how these parameters are estimated in different studies (absolute vs. relative, and semiquantitative vs. quantitative) might have relevant impact. Nevertheless, the combination of fMRI variables (*ReHo* in particular) and CBF or CMRO_2_ led to satisfactory *K*_i_ spatial description (nearly 80% of group-level, 45% of individual-level variance), with marked amelioration of the pattern of the residuals, especially in areas with the strongest positive “outliers” (posterior cingulum). This confirms our hypothesis: the combination of direct hemodynamic and metabolic information, as provided by CBF and CMRO_2_, with information on spontaneous brain activity provided by the BOLD signal explains a large portion of individual-level variance (nearly half) in glucose metabolism (as expressed by *K*_i_). On the other hand, the difficulty in reaching a fully satisfactory description, particularly for *k*_3_ spatial variability (18% at individual level), despite the high number of available features (rs-fMRI, CBF, CMRO_2_), calls for more extensive exploration, potentially looking into other measures, both structural/molecular, such as the spatial distribution of hexokinase isoforms (HK1, HK2), or activity-related, e.g., from electrophysiological signals, which could also help better understand the metabolic role played by *ReHo*, whose biological underpinnings are still unclear.

A separate discussion is warranted for [^18^F]FDG delivery (*K*_1_), which has the most peculiar and interesting spatial distribution, with a markedly posterior pattern (visual cortex, cerebellum, thalamus as “top” parcels), and apparent even in the earliest studies with low-resolution PET cameras^20^. As mentioned in these prior studies^20^, one might be tempted to consider vascular territory effects (i.e., the highest *K*_1_ values largely encompass the posterior circulation), but this does not seem to be a satisfactory explanation, not least because the anterior circulation also provides blood flow to the posterior cerebral territories via the posterior communicating arteries. Notably, the blood volume contribution to the PET signal (*V*_b_), which may differ among vascular territories due to heterogeneous blood velocity, is accounted for during model fitting. Alternatively, some relationship may exist between *K*_1_ and the expression of different isoforms of glucose transporters (GLUT1 and GLUT3, but also SGLT transporters)^14^. Intriguingly, it is also worth noting that although measures of oxygen extraction fraction (OEF) are largely similar throughout the brain, several studies have noted higher OEF in the occipital cortex^63,64^.

When relating *K*_1_ to BOLD measures, it was the only [^18^F]FDG parameter that had significant bivariate associations with most HRF and sFC features. The relatively strong coupling of HRF and large-scale FC network features with *K*_1_ seems to confirm that blood flow and BBB permeability information (of which *K*_1_ is a combination) are important contributors to the rs-fMRI signal^48^, and potentially more linked to large-scale FC than glucose metabolism itself. However, BOLD-based information alone did not provide sufficient explanation of the spatial distribution of *K*_1_ (group-average R^2^ ∼ 0.35, pooled R^2^ ∼ 0.15). Importantly, the inclusion of CMRO_2_ in the fMRI-based models markedly increased explained variance of the individual *K*_1_ data (pooled R^2^ ∼ 0.3), improving the pattern of the residuals. This finding is consistent with previous reports separately describing the posteromedial spatial distributions of [^18^F]FDG *K*_1_^21^ and CMRO_2_ ^62^ (and OEF). Notably, some key differences between the two parameters can also be identified: the cerebellum, in particular, is a “hotspot” for *K*_1_ only; its peculiar structural and physiological characteristics might explain its high [^18^F]FDG delivery, including its different glia-to-neuron ratio^65^, density and type of glucose transporters, different lumped constant^14^, higher *E* and PS product^66^ etc. Of note, CBF was not as effective as a predictor for *K*_1_; this is, however, not surprising, since [^18^F]FDG is not a highly extracted tracer (average *E* < 20%)^66^, and *K*_1_ and CBF are nonlinearly related (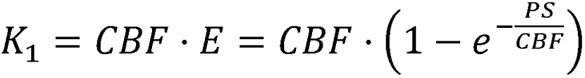 (PS as the permeability-surface product), with *E* having some regional heterogeneity^66^. From a physiological standpoint, the strong association between the delivery of glucose (*K*_1_) and the delivery and consumption of oxygen (CMRO_2_), is interesting and highlights *K*_1_’s metabolic relevance. Further indicators of the metabolic role played by *K*_1_ are the correlations between the strength of its coupling with BOLD and/or CMRO_2_, and peripheral metabolism. Only the *K*_1_-BOLD and *K*_1_-CMRO_2_ coupling strengths (not *K*_i_ and *k*_3_) are related to participant sex, weight and insulin levels, which is consistent with our findings for SUVR^13^. This seems to suggest that individual differences in peripheral metabolism may alter how glucose is delivered to supply the brain’s functional needs^67^, thereby affecting individual *K*_1_-BOLD coupling. This is an interesting area of investigation into the crosstalk between brain and periphery, which could help to reveal how and why metabolic diseases, such as diabetes, lead to increased risk of developing neurological disorders (e.g., Alzheimer’s disease^68^).

Some limitations need to be considered. First, PET and rs-fMRI data were not acquired simultaneously. While this may introduce some within-individual variability^69^, we have shown a better match between SUVR and rs-fMRI using sequentially acquired data in our previous work^13^. This may relate to the high quality of the rs-fMRI data employed in this study (HCP Aging^70^). Second, PET kinetic estimates in this study are not truly fully quantitative. For [^18^F]FDG, using an image-derived input function, which is likely to still be affected by PVEs due to the limited spatial resolution of the HR+ scanner (FWHM ∼5 mm)^71^, may make *K*_i_, *K*_1_ and *k*_3_ estimates biased. However, their *relative* spatial distribution across ROIs, which was the focus of our analyses, is expected to not be impacted. The same reasoning applies to [^15^O]H_2_O and [^15^O]O_2_ data. Third, the low spatial resolution of the HR+ scanner and relatively high noise level in the data makes nonlinear fitting of complex compartment model structures problematic. However, the variational Bayesian framework^31^ we used allowed us to retrieve accurate and precise estimates at the voxel level even in such contexts. We thus believe our [^18^F]FDG kinetic parameter maps are faithful representations of their spatial distribution. Re-assessing these results on high-sensitivity, high-resolution PET scanners^72,73,74^ will be important to confirm the reproducibility of these spatial patterns, possibly capture more details, and further validate their biological meaning. Fourth, it must also be remembered that the BOLD signal is only an indirect measure of neuronal activity, which is subjected to significant contamination from systemic modulations (heart rate variability, vasomotion, respiratory volume variability etc.)^75^. Lastly, despite the use of sophisticated modeling, this work remains correlational: only a controlled perturbational approach may fully elucidate causal links between glucose metabolism and spontaneous activity.

In conclusion, we have comprehensively assessed the physiological information contained in [^18^F]FDG dynamic PET data from a large dataset of 47 healthy individuals, estimating both the macroparameter *K*_i_ (uptake), and the rate constants *K*_1_ (delivery) and *k*_3_ (phosphorylation), with unprecedented spatial detail, to demonstrate how the microparameters add relevant information. We then used the combination of rs-fMRI measures previously identified as the best suited to explain SUVR variance across regions, and verified that they explain a similar portion of *K*_i_ variability (less so for *K*_1_ and *k*_3_). Metrics based on BOLD local properties (*ReHo*) are again most tightly related to glucose metabolism (*K*_i_ and *k*_3_). The inclusion of CBF and CMRO_2_ markedly improves the description of *K*_i_; moreover, *K*_1_ was coupled with CMRO_2_ more than any other feature. Overall, this work enriches the landscape of our understanding of the interplay between PET-and BOLD-derived variables, which reflect complex interactions between brain metabolism (CMRglc, CMRO_2_), blood flow and neural activity. With high-performance PET scanners, assessment of glucose delivery (*K*_1_) and hexokinase activity (*k*_3_) via [^18^F]FDG PET may become useful for evaluating disorders of the brain (e.g., Alzheimer’s disease^19^; traumatic brain injury^21^) and other organs^76^.

## Authors’ contributions

TV and AB designed the research. TV analyzed the data. TV, JJL, AGV, MSG, MC and AB interpreted the results. TV wrote the manuscript. TV, JJL, AGV, MSG, MC and AB revised the manuscript.

