## Supplementary Materials for "The brain’s “dark energy” puzzle *upgraded*: [^18^F]FDG uptake, delivery and phosphorylation, and their coupling with resting-state brain activity"

Tommaso Volpi, Ph.D

801 Howard Ave,

06519 New Haven, CT, USA

+1 2037852385

Alessandra Bertoldo, Ph.D

Via Gradenigo 6/B,

35122 Padova, Italy

+39 0498277694

### Supplementary Methods

*Imaging protocols*

For each participant, a structural MRI scan was performed to provide anatomical information. High-resolution structural images were acquired on a Siemens Magnetom Prisma scanner using a 3D sagittal T1-weighted magnetization-prepared 180° radio-frequency pulses and rapid gradient-echo (MPRAGE) multi-echo sequence (TE = 1.81, 3.6 , 5.39, 7.18 ms, TR = 2,500 ms, TI = 1,000 ms, 0.8×0.8×0.8-mm voxels). The final T1w image was obtained as the average of the first two echoes[1]. Additionally, T2* gradient-echo echo planar imaging (GE-EPI) data were acquired (TR/TE=800/33 ms, flip angle 52°, voxel size 2.4×2.4×2.4 mm, MB 6, 375 volumes for total scan time of 5 min), together with two spin-echo (SE) acquisitions (TR/TE=6000/60 ms, flip angle 90°) with opposite phase encoding directions (AP, PA).

All subjects underwent one [$^{18}$F]FDG PET scan and two sets of three $^{15}$O scans ([$^{15}$O]CO, [$^{15}$O]$H_2O$, and [$^{15}$O]$O_2$). [$^{18}$F]FDG scans were performed on a Siemens model 962 ECAT EXACT HR+ PET scanner (Siemens/CTI)[2], as previously described[3], after i.v. bolus injection of 5.1 ± 0.3 mCi (187.7 ± 12.1 MBq) of [$^{18}$F]FDG. Dynamic acquisition of PET emission data continued for 60 min. [$^{15}$O]$H_2O$ and [$^{15}$O]$O_2$ scans were also performed on the Siemens EXACT HR + scanner (Siemens/CTI)[2], after i.v. bolus injection of 49.6 ± 2.3 mCi (1835.2 ± 85.1 MBq) for [$^{15}$O]$H_2O$, and inhalation in room air of 66.5 ± 6.7 mCi (2460.5 ± 247.9 MBq) for [$^{15}$O]$O_2$. Dynamic acquisition of PET emission data continued for 3 min for both [$^{15}$O]$H_2O$ and [$^{15}$O]$O_2$. Subject head movements during scanning were restricted by a thermoplastic mask. All PET images were acquired in the eyes-closed waking state. No specific instructions were given regarding cognitive activity during scanning other than to remain awake. PET data were reconstructed via filtered back-projection (ramp filter, 5 mm FWHM) as 128x128x63 matrices. Attenuation correction was performed using the participant's own transmission scan. The chosen reconstruction grid for [$^{18}$F]FDG consisted of 52 frames of increasing duration (24 x 5 s frames, 9 x 20 s frames, 10 x 1 min frames, and 9 x 5 min frames), while for [$^{15}$O]$H_2O$ and [$^{15}$O]$O_2$ it consisted of 49 frames (35 x 2 s frames, 6 x 5 s frames, and 8 x 10 s frames). In the case of [$^{18}$F]FDG, venous samples for plasma glucose determination were obtained just before and at the midpoint of the scan to verify that glucose levels were within normal range throughout the study. Also, venous samples were collected to assess [$^{18}$F]FDG plasma concentration, with two possible sampling schedules: for most subjects, sampling occurred 20, 30, 45 minutes after injection of the radiotracer, whereas, for a minority of subjects (n = 10), samples were acquired after 30, 40 and 50 minutes. Each sample consisted of about 2 ml, half of which was used to measure radioactivity in plasma. Radioactivity counter measurements was given in counts per 12 seconds. The counter's efficiency (0.2707 cps/Becquerels) was experimentally determined[4].

*Structural MRI preprocessing*

Structural T1w images were N4 bias field-corrected[5], skull-stripped, and segmented into gray matter (GM), white matter (WM) and cerebrospinal fluid (CSF) using SPM12[6]. T1w images were normalized to the symmetric MNI152 2009c atlas[7] via nonlinear diffeomorphic registration[8]. The Schaefer functional atlas (200 parcels, 17 networks)[9] was registered to T1w space by inverting the obtained nonlinear transformation. The Schaefer ROIs were supplemented by 16 subcortical ROIs from the Hammers atlas[10] (bilateral hippocampus, amygdala, caudate, accumbens, putamen, pallidum, thalamus, cerebellum).

*[$^{18}$F]FDG PET kinetic modeling*

Dynamic PET data were motion-corrected using in-house implementations of PMOD (www.pmod.com) and FSL's *mcflirt*[11]. A static PET image was obtained by summing late PET frames (40-60 min) after motion correction. To perform PET kinetic modeling, an image-derived input function (IDIF) was extracted from dynamic PET data using a semi-automatic pipeline[12], including 1) segmentation of the internal carotid arteries from a pseudo-angiography image using a *vesselness* algorithm[13], 2) selection of "hot voxels" according to peak amplitude and time-to-peak, 3) parametric clustering[14] (k-means algorithm, k=2, squared Euclidean distance, 500 replicates) on seven parameters calculated on the TAC of each voxel, with the cluster having the highest peak centroid being selected and used to derive the raw IDIF, 4) IDIF model fitting using a modified version of Feng's model[15], 5) Chen's spillover correction[16] using three venous samples (obtained after 20 min post-injection), 6) IDIF shift correction, to correct for delay between the carotids and the voxel of interest.

Voxel-wise estimation of Sokoloff's model parameters was performed using a Variational Bayesian approach[17], according to the following pipeline:

- a k-means clustering approach is applied to dynamic PET data, extracting 6 GM and 5 WM clusters (as from the tissue segmentations linearly mapped to PET space),

- nonlinear fitting of Sokoloff's model (using weighted nonlinear least squares, with weights chosen as the inverse of the variance of the PET measurement error[18]) is performed at the region level, i.e., on the 11 cluster centroids,
- voxel-wise estimates of the model parameters are obtained via Variational Bayesian inference using prior distributions derived from cluster-wise estimates.

We refer to [17] for more details on the estimation procedure.

Parametric maps (i.e., at voxel level) of $K_1$ [mL/cm$^3$/min] (tracer inflow), $k_2$ [min$^{-1}$] (efflux), $k_3$ [min$^{-1}$] (phosphorylation), $V_b$ [%] (blood volume fraction) were obtained for each individual. The parametric map of $K_i$ [mL/cm$^3$/min] (irreversible tracer uptake) was obtained by the solving $K_i = \frac{K_1 k_3}{k_2+k_3}$ at voxel level.

The group average maps of $K_i$, $K_1$, $k_3$ are reported in **Figure 2 A, B, C**, respectively, while $k_2$ and $V_b$ are reported in **Figure S1**.

The $K_i$, $K_1$, $k_3$ parametric maps were parceled at the individual level with the Schaefer + Hammers atlas: ROI-level parameter estimates were extracted from the Schaefer and Hammers parcels, which had been linearly mapped from T1w to PET space, by averaging over voxels within the GM segmentation (probability > 0.8 of belonging to GM to obtain a seamless hard labeling).

Importantly, the GM segmentation provided by SPM, being quite conservative, allows to extract an average TAC which is as free of partial volume effects (PVEs) as possible[19]. Moreover, spatial smoothing of the PET data during processing was avoided, further minimizing PVEs.

The region-wise $K_i$, $K_1$, $k_3$ values were within-individual normalized via z-scoring, i.e., centered with respect to their mean and divided by the standard deviation across ROIs.

*[$^{15}$O]H$_2$O and [$^{15}$O]O$_2$ PET kinetic modeling*

The differential equation of the [$^{15}$O]H$_2$O tracer's one-tissue compartment model

$$\dot{C}_1(t) = K_1 C_\mathrm{p}(t) - k_2 C_1(t)$$

with $C_1$ as the tissue tracer concentration and $C_\mathrm{p}$ as the arterial input, was linearized as follows:

$$C_1(t) = K_1 \int_0^t C_\mathrm{p}(\tau) d\tau - k_2 \int_0^t C_1(\tau) d\tau$$

to identify the $K_1$ [mL/cm$^3$/min] (inflow), which in the case of [$^{15}$O]H$_2$O corresponds to the CBF, and $k_2$ [min$^{-1}$] (efflux).

Since arterial samples to determine $C_\mathrm{p}(t)$ were not available, and the data were too noisy to extract an IDIF from the carotid signals, we used a model-based IDIF approach similar to [20], which reconstructs the $C_\mathrm{p}$ by rearranging equation 4 as follows:

$$C_\mathrm{p}(t) = \frac{1}{CBF^{WB}} \dot{C}_1(t) + \frac{k_2^{WB}}{CBF^{WB}} C_1(t)$$

With $C_1(t)$ which here corresponds to the whole-brain average tissue TAC from dynamic [$^{15}$O]H$_2$O data, $CBF^{WB}$ as a whole-brain average CBF value, and $\frac{k_2^{WB}}{CBF^{WB}}$ corresponding to $\frac{1}{\lambda}$ ($\lambda$ is the blood-brain partition coefficient for water). $CBF^{WB}$ and $\lambda$ are chosen *a priori* as 0.5 mL/cm$^3$/min and 0.9 ml/cm$^3$, respectively, based on population values. The raw IDIF curve was fit with a Gamma-variate function as in [14] to regularize its shape.

Since $K_1$ is directly dependent on the amplitude of the $C_\mathrm{p}(t)$, the final average CBF value will be approximately close to the chosen value for whole-brain $K_1$. This makes the result of this approach a *relative* CBF map. However, for this work, we do not need *absolute* estimates of CBF as we only aim to compare relative spatial distributions between [$^{18}$F]FDG parameters, fMRI variables, and CBF and $CMRO_2$.

To estimate $CMRO_2$, a reference-tissue modeling approach based on [21] was employed. Voxel-wise $CMRO_2$ values were obtained via the following equation:

$$CMRO_{2i} = CMRO_2^{WB} \frac{\int_0^T C_{1i}(t)dt + \frac{CBF_i}{\lambda}\int_0^T \int_0^t C_{1i}(u)dudt}{\int_0^T C_1^{WB}(t)dt + \frac{CBF^{WB}}{\lambda}\int_0^T \int_0^t C_1^{WB}(u)dudt} \quad \text{for } i = 1, \dots p \text{ voxels}$$

with $CMRO_2^{WB}$ as whole-brain average $CMRO_2$ value, $C_{1i}$ as the voxel-wise [$^{15}$O]$O_2$ tissue TAC, $CBF_i$ as the voxel-wise CBF values, obtained from [$^{15}$O]$H_2O$ PET modeling, $C_1^{WB}$ as the whole-brain [$^{15}$O]$O_2$ tissue TAC. The $CMRO_2^{WB}$ value was obtained by

$$CMRO_2^{WB} = C_a^{O_2}\ CBF^{WB}\ \frac{\left(S_a^{O_2} - S_v^{O_2}\right)}{S_a^{O_2}}$$

with $C_a^{O_2}$ as the $O_2$ arterial tension, set to the literature value of 90 mmHg, and $S_a^{O_2}$ as the $O_2$ arterial saturation, set to 98%. Due to the use of literature values for $C_a^{O_2}$, $S_a^{O_2}$, $CBF^{WB}$, the result of this approach is a *relative* $CMRO_2$ map as well.

The group-average maps of CBF and $CMRO_2$ are reported in **Figure S1**.

The CBF, $CMRO_2$ parametric maps were parceled at the individual level with the Schaefer + Hammers atlas (GM-masked to minimize PVEs). Spatial smoothing of the PET data was avoided. The region-wise CBF, $CMRO_2$ values were within-individual normalized via z-scoring, i.e., centered with respect to their mean and divided by the standard deviation across ROIs.

*Functional MRI data pre-processing*

The fMRI data were analyzed using an in-house pipeline based on the Human Connectome Project minimal preprocessing pipeline[22]. The first four fMRI volumes were discarded to avoid non-equilibrium magnetization effects. The remaining volumes were corrected for slice timing differences[23] and magnetic field distortion[24], and realigned to the median volume using FSL's *mcflirt*[25]. A template EPI volume was obtained from realigned fMRI data with *antsBuildTemplate*[8] and used to estimate an affine transform (*flirt*, FSL), subsequently employed to map main tissue segmentations obtained from the T1w image to the EPI space.

Nuisance signals, including motion parameters and their first order derivatives, and the first 5 temporal principal components obtained after principal component analysis of WM and CSF EPI signals[26], were regressed out from all brain voxels in native EPI space[27]. Finally, the BOLD signal was high-pass filtered with a cut-off of 0.008 Hz. No low-pass filter was applied, as the higher frequency components (0.1-0.25 Hz) of BOLD are likely to provide relevant neural information[28].

Motion correction was adapted to the features to be extracted. For features where it was important to preserve the temporal structure of the BOLD signal (e.g., time-varying FC, time-varying *ReHo*, HRF-based measures), motion-corrupted volumes were corrected by *despiking* with a cubic and spline interpolation, using the *icatb_despike_tc* function from the Group ICA Toolbox GIFT[29], in order to avoid extreme censoring methods that would interrupt the temporal autocorrelation structure of the data[30]. For features that were more robust to *censoring* (e.g., static FC, *ReHo*), motion-corrupted volumes with frame-wise displacement (FD) higher than 0.3 mm were discarded before sFC calculation[31]. Mean FD and the number of censored volumes were evaluated for every participant, to ensure that a sufficient number of viable frames was available.

Pre-processed EPI signals were parceled within each region from the Schaefer + Hammers atlas, which had been linearly mapped from T1w to EPI space, by averaging over voxels within the GM segmentation (probability > 0.8 of belonging to GM).

*Resting-state fMRI feature extraction*

Feature extraction as well as subsequent analyses were performed in MATLAB (The Mathworks, Natick, MA). Fifty distinct features were obtained from the rs-fMRI signal, either at the voxel level or at the parcel level, as in our previous work[32]. The extracted features were chosen as descriptors of different aspects of the BOLD 1) *signal*, 2) *HRF*, 3) *sFC*, and 4) *tvFC*. A list of the features and their acronyms is reported in **Table 1**.

1. *Signal and local features*

The temporal *median*, median absolute deviation (*MAD*) and *skewness* of the parcel-wise BOLD signal, i.e., the first, second, and third-moment statistics of the BOLD time series distribution, were calculated. The average BOLD signal has been found to be related not only to activity but also cell density[33,34], and the BOLD signal variability is known to carry significant physiological information on cellular properties[35,36] and also to be a correlate of cerebrovascular reactivity[37]; the skewness, which captures extreme BOLD events, has been found to be related to structural connectivity (SC), with more connected regions exhibiting resting-state activity with high negative skewness[38]. Moreover, nonlinear metrics of BOLD signal *complexity* were computed, in particular *approximate entropy* (ApEn)[39] and *range approximate entropy* (rApEn)[40].

ApEn quantifies the mean negative log-probability that an $m$-dimensional state vector will repeat itself at dimension ($m + 1$). It is recommended that the tolerance is corrected as $r \times SD$ ($SD$ being the standard deviation of the signal) to account for amplitude variations across different signals. The embedding dimension $m$ was set equal to 2, and the tolerance $r$ was set to 0.2 multiplied by the SD of the signal[39]. rApEn[40] is more robust to nonstationary signal amplitude changes and sensitive to changes in the Hurst exponent with respect to ApEn, thus being more appropriate to evaluate self-similarity in the signal. For what concerns the parameter $r$, it is constrained to the range [0,1], so it does not need any amplitude correction. A first-order autoregressive AR(1) model was also fit to the windowed BOLD time series by minimizing the forward prediction error in the least squares sense; the Yule-Walker equations were solved by the Levinson-Durbin recursion, obtaining the AR(1) reflection coefficients, whose absolute value was taken as the time dependence between $y(n)$ and $y(n-1)$. The *exponents* of a first-order autoregressive AR(1) model fit to the BOLD signal are tightly linked to the Hurst exponent, that had previously been found to have physiological and cognitive relevance[40].

At the voxel level, instead, the BOLD signal's spectral content was quantified by the amplitude of low frequency fluctuations (*ALFF*), as signal power within [0.01, 0.1] Hz band[41]. The local coherence of the BOLD signal was described by "regional homogeneity" (*ReHo*)[42], computed as Kendall's W coefficient of concordance among one voxel's time series and its 27 neighbors'. Parcel-wise ReHo and ALFF values were then extracted as the average of the voxels within the region (GM-masked). *Time-varying ReHo* was computed with a sliding window approach (window size: 30 TRs, step: 1 TR), as Kendall's coefficient of concordance amongst neighboring voxels within each time window[43]. Motion-corrupted volumes were corrected by voxel-wise despiking with a cubic and spline interpolation, using the *icatb_despike_tc* function from the

Group ICA Toolbox GIFT[29], in order to avoid interrupting the temporal autocorrelation of the data[30]. ReHo time courses were extracted at the parcel level by averaging voxels within a region. Regional ReHo variability was calculated as nonparametric median absolute deviation (*MAD*) and coefficient of variation (*CV%*) of the parcel-wise time series, i.e., $CV_{nonpar} = \frac{MAD}{median} \cdot 100$

2. *HRF features*

The parcel-wise BOLD signal was subjected to a blind deconvolution algorithm[44,45] employing the *rs-HRF toolbox v2.0* (*https://www.nitrc.org/projects/rshrf*). This approach builds on previous work describing the rs-fMRI signal as a point process, with events that exceed a given threshold governing the dynamics. In a linear, time-invariant (LTI) system framework, the BOLD signal, y(t), is thus modeled as the convolution of the HRF, h(t), and the underlying neural states, s(t), with the addition of an error term, e(t):

$$y(t) = s(t) \otimes h(t) + e(t)$$

Before deconvolution, the high-pass filtered BOLD signal was despiked using a hyperbolic tangent squashing function. The HRF was estimated as the linear combination of basis vectors for a smooth finite impulse response (sFIR)[45,46], which captures any shape up to a given smooth frequency filter, without constraining the HRF to the canonical model (two gamma functions with time and dispersion derivatives[47]). BOLD pseudo-events were detected using a threshold, which was set to the default value of 1 SD from the mean of the BOLD signal, and their parcel-wise number was calculated (*peaks-BOLD*). Serial correlations in BOLD time series due to aliasing of biorhythms and unmodelled neuronal activity were accounted for using an autoregressive AR(1) model during parameter estimation[44]. The outputs of the deconvolution process were:

- three parcel-wise HRF parameters, i.e., 1) *HRF height*, which was used for further analyses; 2) HRF full-width-at-half-maximum (FWHM); 3) HRF time-to-peak;
- the time course of the parcel-wise HRFs (16 time points, with time bins of 2 seconds, each corresponding to one TR);
- the time course of the deconvolved BOLD signal.

For each subject, a pairwise Pearson's correlation matrix was calculated from the parcel-wise HRFs, as the matrix of zero-lag temporal correlations between HRF time series of each pair of regions, interpreted as signals of (putatively) vascular origin. These subject-wise "HRF connectivity" matrices were Fisher r-to-z transformed, and then thresholded retaining only connections associated with weights over a pre-defined connection density, set to the 80$^{th}$ percentile[48].

The following nodal topological features were estimated from these matrices using the Brain Connectivity Toolbox[49].

Node degree (DEG), defined as the number of links connected to a node, was used to characterize network structure and local connectivity. Node strength (STR), i.e., the sum of all link weights, was used to complement node DEG as a measure of connectivity profile. Eigenvector centrality (EC), which uses eigendecomposition of the HRF correlation matrix to measure if strong connections tend to link nodes with equally strong connections, accounting for the importance of indirect pathways[50], was calculated as a node centrality measure. Betweenness centrality (BC), another node centrality measure, was calculated as the number of shortest paths between nodes passing through a specific node[51]. Network segregation was measured by the clustering coefficient (CC), which locally represents the number of triangles around an individual node over the number of connected triples in the network[52,53]. Segregation was also evaluated through local efficiency

(LE), i.e., the ratio of the number of connections between a node's neighbors to the total number of possible links[54]. Finally, another measure of integration was the global efficiency (GE), which is the inverse shortest path length in the network[54].

3. *Static FC features*

sFC matrices were obtained as pairwise Pearson's correlation coefficients of the BOLD time series across brain regions, which were subsequently Fisher r-to-z transformed. Motion-corrupted volumes with frame-wise displacement (FD) higher than 0.3 mm were discarded before sFC calculation[31]. Subject-wise sFC matrices were thresholded ($80^{th}$ percentile[48]). Topological features of sFC matrices (node DEG, STR, EC, BC, CC, LE, GE) were estimated using the Brain Connectivity Toolbox[49], with the aforementioned purposes.

In addition to the more frequently employed '*magnitude* FC' approach, we also characterized FC as BOLD *phase* coherence, employing the Leading Eigenvector Dynamic Analysis (LEiDA) framework[55,56]. After demeaning and detrending the BOLD time series, the parcel-wise BOLD signal phases, $\theta(n,t)$, were estimated using the Hilbert transform, which expresses a given signal as $x(t) = A(t) \cdot \cos\theta(t)$, where $A(t)$ is the time-varying *amplitude* and $\theta$ is the time-varying *phase*. BOLD phase coherence is then calculated at each single time point *t*, which results in a 3D matrix of time-varying phase-locking values (tvPLV) between each pair of brain areas *n* and *p* at each time *t*, estimated as:

$$\text{tvPLV}(n,p,t) = \cos\left(\theta(n,t) - \theta(p,t)\right)$$

By using the cosine function, two regions with no BOLD phase difference at a given TR will have a $\text{tvPLV}(n,p,t) = \cos(0°) = 1$, while time points when BOLD signals have 180° phase difference have a $\text{tvPLV}(n,p,t) = \cos(180°) = -1$, which makes the tvPLV(t) matrix symmetric with values ranging between -1 and 1. Then, the leading eigenvector (LEig) of the tvPLV(t) matrix at time t is computed to capture the main orientation of regional BOLD phases over all other brain areas. The *signs* of LEig, when both positive and negative, naturally divide brain regions into two clusters according to their BOLD phase relationship, while the *magnitude* of each element indicate the strength with which each brain region belong to its community. It is noteworthy that the LEig consistently represents > 50% of the variance in phase coherence at all time points.

The parcel-wise median value of LEig across time was calculated in every subject and interpreted as a 'static' measure of phase coherence (*med-LEig*).

#### 4. *Time-varying FC features*

tvFC was computed with a sliding window approach (window size: 30 TRs, step: 1 TR), as Fisher r-to-z transformed Pearson's correlation.

Motion-corrupted rs-fMRI volumes were dealt with by parcel-wise despiking with cubic and spline interpolation, using the *icatb_despike_tc* function from the Group ICA Toolbox GIFT[29]: despiking is frequently used in tvFC studies (e.g., [57,58]), in order to avoid more extreme censoring methods that would interrupt the temporal autocorrelation structure of the data and result in sliding windows of differing lengths. Sliding windows were thresholded using the connection density threshold approach ($80^{th}$ percentile): FC weights were selected on the population sFC matrix, and then propagated to the single sliding windows, to assess the temporal variability of the connections that are most likely to be significant at the population level. The same nodal graph metrics used in the sFC analysis (STR, CC, BC, EC, LE, GE) were computed for each window in every subject. Three metrics to quantify temporal variability across sliding windows were selected and applied to the graph metrics' time series at the parcel level:

a) nonparametric coefficient of variation (*CV%*, MAD/median ratio), as a measure of fluctuation of the graph metric around its average value[59];

b) temporal median of the absolute value of 1st order differentials (*mdiff*) between graph metrics' values in adjacent sliding windows divided by the absolute value of the previous window, as a measure of average deviation of the graph metric's value $x_{it}$ from its previous values $x_{it-1}$:

$$\mathrm{mdiff}_{\mathrm{x_i}} = \mathrm{median}\left(\frac{|x_{it} - x_{it-1}|}{|x_{it-1}|}\right)$$

c) *sample entropy* of graph metrics' time series as a measure of graph metrics' time series complexity[60].

In addition to the *magnitude* tvFC metrics, the regional *MAD*, *CV%* and *mdiff* of the LEigs were calculated as metrics of temporal variability of *phase* coherence.

*Bivariate associations between [$^{18}$F]FDG kinetic parameters and functional features*

The bivariate relationship between node-wise [$^{18}$F]FDG kinetic parameters ($K_i$, $K_1$ and $k_3$) and rs-fMRI properties or CBF/$CMRO_2$ was assessed at the group level (naïve average data, NAD approach), employing the region-wise mean values across individuals for [$^{18}$F]FDG kinetic parameters and for each of the 50 extracted features. As most rs-fMRI properties were normally distributed ($p > 0.05$, Shapiro-Wilk test[61]), the association between fMRI features and metabolism across nodes was tested via Pearson's bivariate correlation (significance level 0.05).

As in our previous work[32], Pearson's correlations between [$^{18}$F]FDG kinetic parameters and functional features were tested across regions selected according to linearly increasing percentiles (from 1st to 85th) and decreasing percentiles (from 100th to 15th) of the $K_i$, $K_1$ and $k_3$ distribution. Due to the nonlinearities that emerged from evaluating clusters of nodes, as in our previous work[32], the relationship between [$^{18}$F]FDG kinetic parameters ($par_{ik}$, for $i$ = 1, …, 216 regions, and $k$ = $K_i$, $K_1$ or $k_3$) and each of the 50 rs-fMRI properties ($fMRI_{ip}$, for $p$ = 1, …, 50 features) was also tested with three different bivariate models:

1) a *linear* model,

$$par_{ik} = \alpha_p + \beta_p \cdot fMRI_{ip}$$

2) a *mono-exponential* model,

$$par_{ik} = \alpha_p \cdot e^{\beta_p \cdot fMRI_{ip}}$$

3) a *power law* model,

$$par_{ik} = \alpha_p \cdot fMRI_{ip}^{\beta_p}$$

Model selection was performed according to the residual sum of squares (RSS)[62] to evaluate whether the [$^{18}$F]FDG-fMRI association was better described by a linear or a nonlinear model for each of the 50 features. The RSS values of the nonlinear models (*exp, power*) were expressed in terms of percent difference with respect to the linear model (*lin*), as follows,

$$\Delta_{RSS_1} = \frac{RSS_{lin} - RSS_{power}}{RSS_{lin}} \cdot 100$$

$$\Delta_{RSS_2} = \frac{RSS_{lin} - RSS_{exp}}{RSS_{lin}} \cdot 100$$

in order to perform a ranking. It must be noted that the number of model parameters was equal for the four model structures that were examined (i.e., two, intercept/amplitude $\alpha$ and slope $\beta$).

*Multivariable modeling of the functional-metabolic relationship at group level*

The bivariate model selection process favored the *log-linear* model for multiple regression modeling. At the NAD level, a multiple linear regression approach was employed to assess how much of the group-wise $K_i$, $K_1$ and $k_3$ variance across regions could be explained by the linear combination of different fMRI-based features (plus CBF and $CMRO_2$). The procedure was applied to $K_i$, $K_1$ and $k_3$ separately. The ordinary least squares (OLS) problem was formulated as follows:

$$y = X\beta + \varepsilon$$

where $y$ and $\varepsilon$ are $n \times 1$ vectors of the response/dependent variable (i.e., $K_i$, $K_1$ or $k_3$) and the model error, and $X \in \mathbb{R}^{n \times p}$ is the design matrix of $p$ regressors (i.e., log-transformed rs-fMRI predictors), or design matrix. Before performing OLS regression, all predictors were z-scored, i.e., centered and scaled by their standard deviation (SD) across brain regions. The outcome variable was z-scored as well, eliminating the model intercept. The solution to the OLS problem was formulated as

$$\hat{\beta} = (X^T X)^{-1} X^T y$$

The model was formulated as follows:

$$par_{ik} = \beta_1 \cdot \log fMRI_{i1} + \beta_2 \cdot \log fMRI_{i2} + \cdots + \beta_p \cdot \log fMRI_{ip} + \varepsilon_i$$

for each observation $i = 1, \ldots, n$. The relationships amongst the predictors were evaluated by Pearson's correlation, to assess the presence of strong correlations (i.e., multicollinearity). High multicollinearity amongst predictors is known to result in lower precision, switched signs of the coefficients, and a lack of statistical significance of the multivariable model[63]. Consequently, we examined the condition number of the design matrix,

$$\kappa(X) = \frac{\sigma_{\max}(X)}{\sigma_{\min}(X)}$$

with $\sigma_{\max}(X)$ and $\sigma_{\min}(X)$ as the highest and lowest singular values of *X,* respectively. As a rule of thumb, $\kappa(X)$ requires attention if higher than 30[63]. Moreover, we calculated the variance inflation factors (VIFs) to assess how much each individual predictor contributed to the multicollinearity of the final model[63]. It is well-known that, in the case of overparameterized linear models, OLS is generally not useful, as many standard errors (%SE, i.e., percent standard error divided by the absolute value of the parameter estimates) are too high (%SE > 100%) and the model is not *a posteriori* identifiable, so it should be rejected[64].

The obtained results were evaluated[62] in terms of:

- number of selected features;

- condition number $\kappa(X)$ of the design matrix after selection;

- Pearson's coefficient of determination (ordinary $R^2$);

- BIC, which proves useful in cases with different numbers of parameters;

- RSS, as indicator of goodness of fit;

- parameter %SEs as indicators of the precision of the estimates;

- $\beta$ estimates and their signs, i.e., the concordance with the signs of Pearson's correlation of the predictors with $K_i$, $K_1$ or $k_3$.

*Full hierarchical modeling of the functional-metabolic relationship*

A NAD approach like the one described so far is statistically sound and unbiased only in case of low between-individual variability (BIV). Consequently, a multilevel population modeling approach (mixed-effect model) was employed to characterize in a single stage both the group-level (fixed) and individual-level (random) effects[73] contributing to the relationship between the selected predictor variables and $K_i$, $K_1$ or $k_3$. First, the link between model and $K_i$, $K_1$ or $k_3$ was described at *individual* level by the following equations:

$$y_i = F_i(X_i, \psi_i)$$

$$z_i = y_i + v_i$$

with $y_i$ as the [$^{18}$F]FDG parameter model prediction for the $i^{th}$ individual ($i = 1, \dots, m$), which is a function of $X_i$ (the fixed-effects design matrix composed by the features of individual $i$ – rs-fMRI and/or CBF, $CMRO_2$), and the parameters $\psi_i$ to be estimated for individual $i$; $z_i$ is the vector of the measured data ([$^{18}$F]FDG parameter estimates) of individual $i$ and $v_i$ is the *within-individual variability*, or residual unexplained variability, assumed to be normally distributed with zero mean and variance $\sigma_i^2$.

Second, at *population* level, $\psi_i$ was described by a function combining population parameters (or fixed effects, $\theta$), and the random variability of individual parameters around the population mean (or random effects, $\eta_i$), according to the following assumptions:

$$\eta_i \sim N(0, \Omega)$$

$$\psi_i = \theta + \eta_i$$

where $\eta_i$ is normally distributed with zero mean, independent across individuals and with covariance matrix $\Omega$. Consequently, $\psi_i$ is normally distribution as well. The matrix $\Omega$ was assumed to be full.

The *intra-individual* (first-level) model structure was composed by the nine features selected with the NAD approach, here at individual level. Data normalization was performed within individuals via z-scoring across regions. The *inter-individual* model (second-level) describing the BIV of the parameters was set according to the aforementioned assumptions.

The normality of model residuals $v_i$ (or a reasonable approximation thereof) was assessed at each level by inspecting their histograms, boxplots, and Q-Q plots. The normality of the random effects $\eta_i$ was inspected with histograms and boxplots. Estimation requires solving a penalized least squares problem, which can be optimized via a restricted maximum likelihood approach. The standard errors (SE) were calculated for each $\theta$ parameter estimate as the square root of the diagonal of their covariance matrix. The overall and individual multilevel model $R^2$ were also evaluated. The residual unexplained variability $v_i$ was evaluated by calculating its mean and variability (percent SD divided by mean) across individuals.

### Supplementary Results

*Between-individual variability of [$^{18}$F]FDG kinetic parameters*

Percent variability (CV%) of [$^{18}$F]FDG parameter absolute values ($K_1$, $k_3$, $K_i$) across individuals ($n = 47$) for each brain region is reported in **Figure S2**. Overall, relatively low interindividual variability is apparent for all parameters (< 30% in most regions).

*Correlations between [$^{18}$F]FDG kinetic parameters*

The across-*region* correlations (Pearson's r) between group-average [$^{18}$F]FDG kinetic parameters are reported in **Figure S3**. Of note, $k_2$ and $k_3$ are negatively correlated ($r = -0.58$, $p < 10^{-5}$). The ratio between $K_i$ and $K_1$ ($= k_3/(k_2+k_3)$), which represents the probability that each tracer molecule is trapped/phosphorylated and is expected to be a good approximation of $k_3$, was indeed very similar to $k_3$ ($r = 0.94$, $R^2 = 0.89$). We also assessed the across-*individual* associations (Pearson's r, $R^2$) among pairs of [$^{18}$F]FDG parameters region by region (**Figure S5**). The correlations of $K_i$ with the microparameters are moderate to high, both for $K_i$ vs. $K_1$ (mean ± SD of absolute r values: 0.561 ± 0.089) and $K_i$ vs. $k_3$ (0.636 ± 0.078). The microparameters, instead, are overall uncorrelated ($K_1$ vs. $k_3$: 0.128 ± 0.109).

*Nonlinear associations between [$^{18}$F]FDG parameters and functional features*

$K_i$ shows strong and significant correlations mainly in the left portion of the matrix (**Figure S8**, *left panel*, linearly decreasing percentiles of $K_i$, i.e., after removing more and more high $K_i$ nodes). A similar, although weaker, pattern of correlations emerges for $k_3$ (**Figure S8**, *right panel*), while $K_1$ is enriched by significant correlations with *ALFF* and CBF-related features (*MAD-BOLD*, *peak-*

*HRF*) in the high-$K_1$ area, i.e., on the right of the $K_1$ matrix (**Figure S8**, *middle panel*), as well as with the other HRF features.

We performed model selection to assess whether a linear, exponential, or power law model would better describe the bivariate spatial relationships between group-average [$^{18}$F]FDG kinetic parameters and rs-fMRI (**Figure S9**). The model selection procedure was performed by evaluating the percentualized differences in RSS between the linear model and both the power ($\Delta RSS_1$) and exponential model ($\Delta RSS_2$). The positive $\Delta RSS_1$ and $\Delta RSS_2$ values (%) are shown in **Figure S9A** for $K_i$ (left), $K_1$ (middle), and $k_3$ (right): the nonlinear models describe the data better than the linear in 48% of the cases for $K_i$, 56% for $K_1$, and 54% for $k_3$ (**Figure S9B**). This confirms a tendency towards nonlinearity in the [$^{18}$F]FDG vs. rs-fMRI bivariate associations in around half of the features, with most strong nonlinear (power law) associations from the sFC and tvFC pools. For this reason, we employed a nonlinear transformation (= natural logarithm) of all the features, as in our previous work[32].

*Group-average correlations of CBF and $CMRO_2$ with rs-fMRI features*

The group-average spatial correlations of CBF and $CMRO_2$ with the rs-fMRI features are shown in **Figure S11.** Of note, $CMRO_2$ is significantly correlated with average rs-fMRI signal of each region (*med-BOLD*), while [$^{18}$F]FDG kinetic parameters are not (**Figure 4**). Also, *ReHo* is only weakly correlated with $CMRO_2$. Moreover, moderate significant correlations are present between $CMRO_2$ and *MAD-BOLD*, *peak-HRF*, both blood flow-related indices. Overall, correlations with the HRF and sFC pool are significant and stronger than for [$^{18}$F]FDG $K_i$ and $k_3$, while a similar pattern is present for tvFC. Notably, the $CMRO_2$-fMRI correlation values have a similar range to those of $K_1$ (0.312 ± 0.058).

### Supplementary Figures

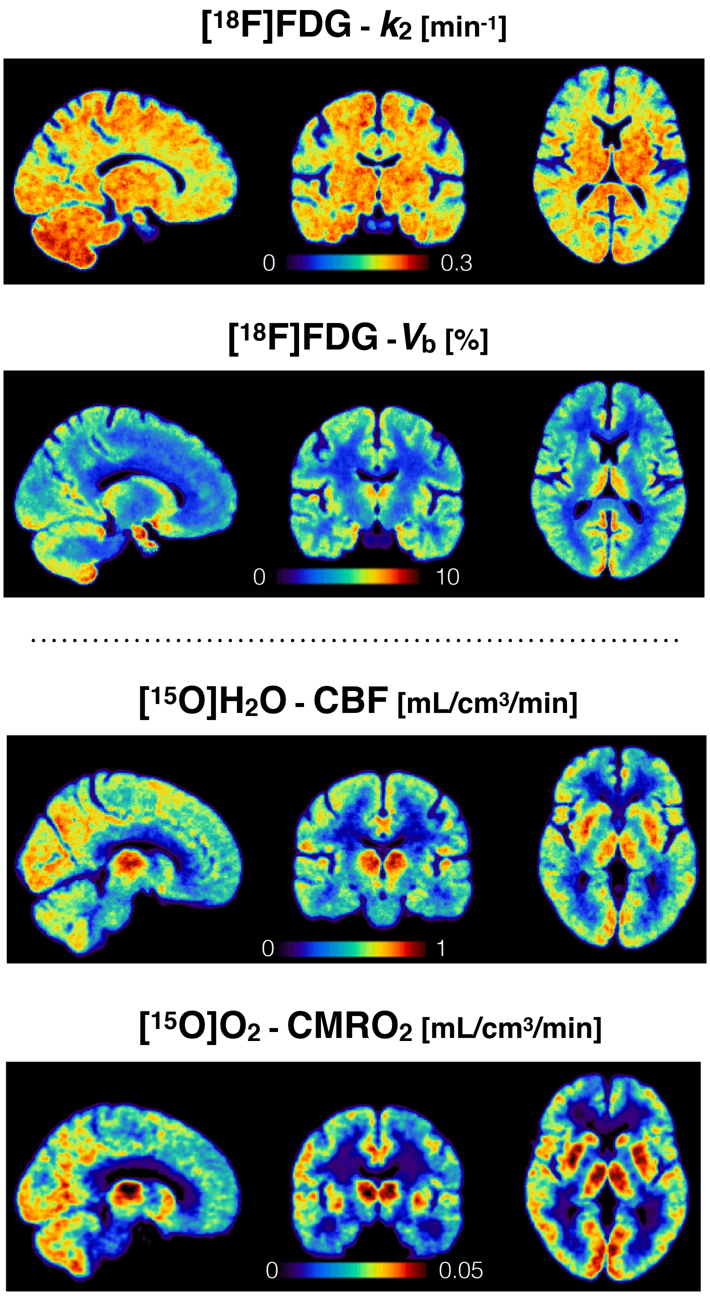


**Figure S1**: Group-average PET parametric maps (n = 47) for [$^{18}$F]FDG-derived $k_2$ and $V_b$, [$^{15}$O]$H_2O$-derived CBF, [$^{15}$O]$O_2$ -derived $CMRO_2$, in MNI space.

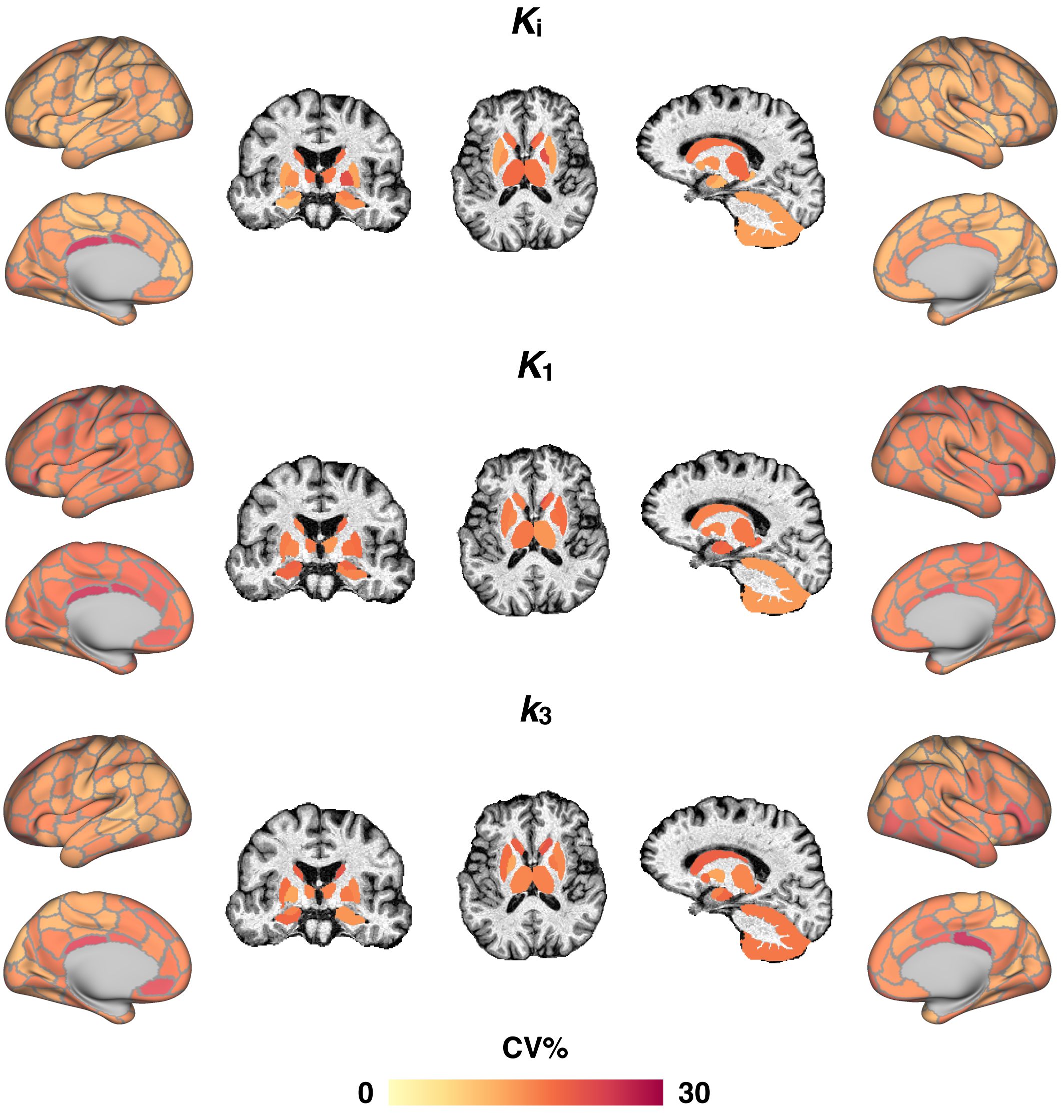


**Figure S2:** Percent variability (CV%) of [$^{18}$F]FDG parameters ($K_i$, $K_1$, $k_3$) across individuals (n = 47) for each brain region.

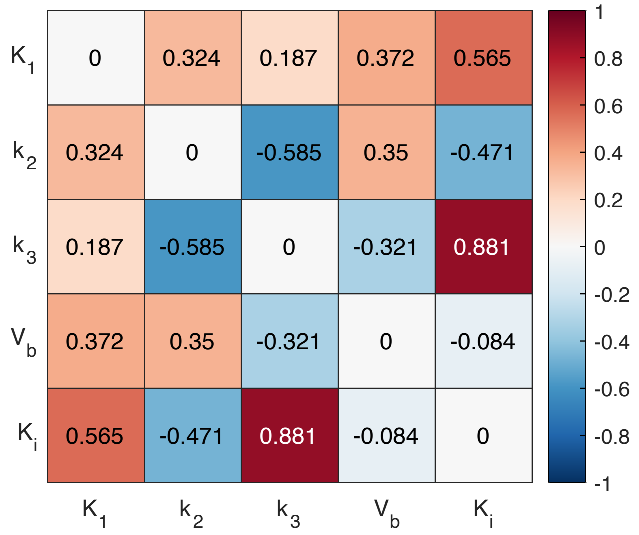


**Figure S3:** Across-region Pearson's correlations between all pairs of [$^{18}$F]FDG kinetic parameters ($K_i$, $K_1$, $k_3$, $V_b$, $K_i$) at group-average level.

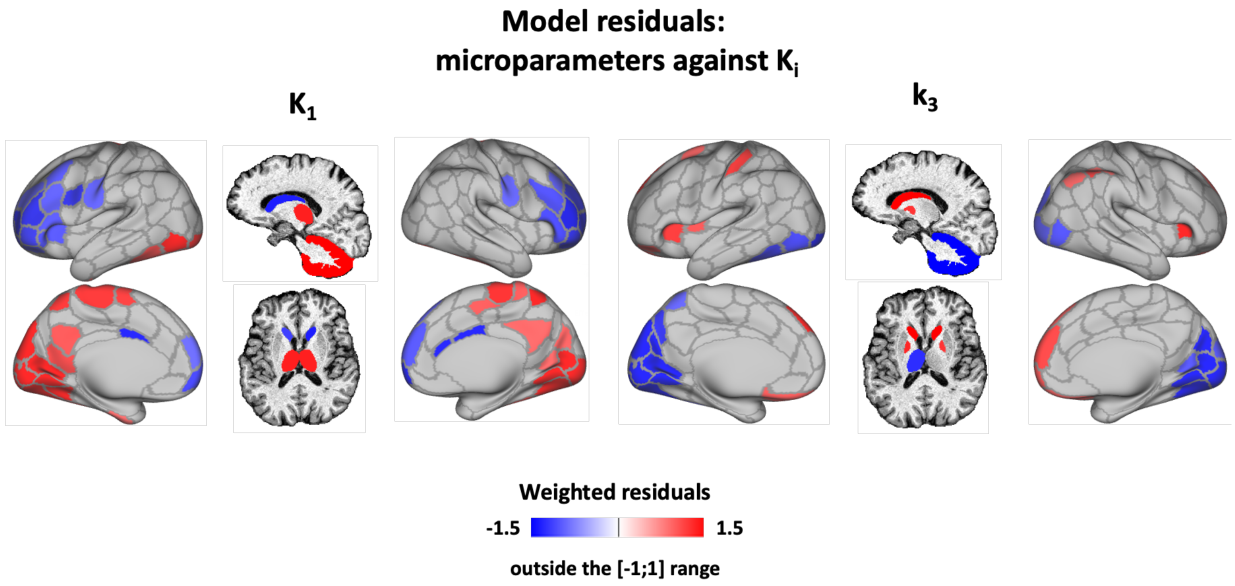


**Figure S4**: Weighted residuals of the linear regression of group-average $K_1$ (left) and $k_3$ (right) on $K_i$; weighted residual values in the [-1;1] range are set to zero.

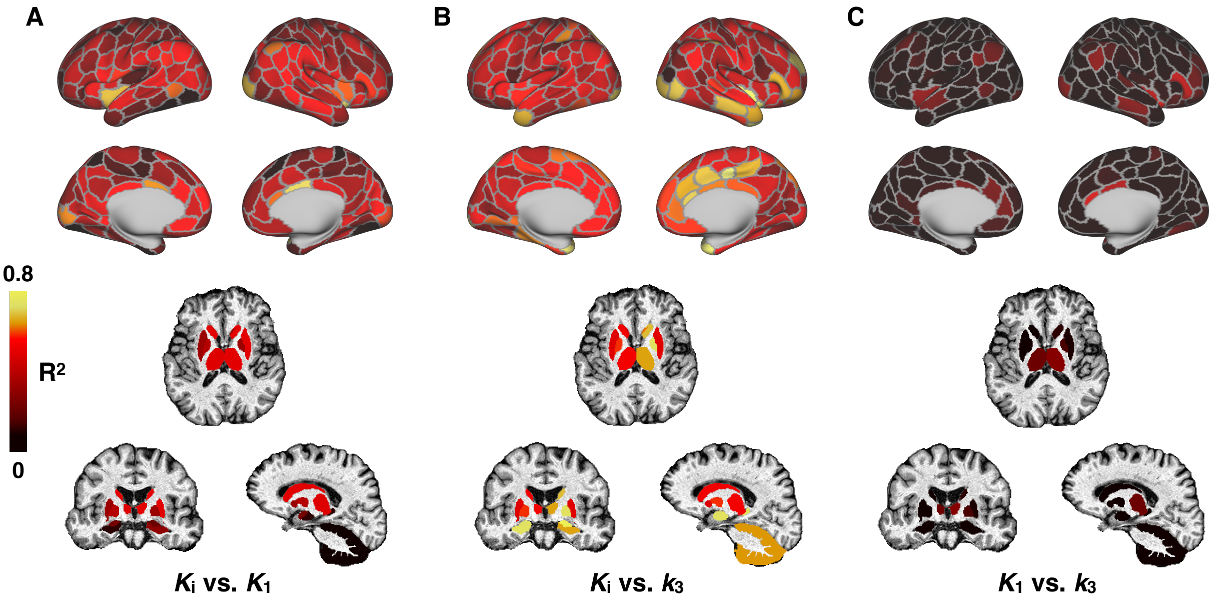


**Figure S5:** Across-individual Pearson's $R^2$ between [$^{18}$F]FDG parameters ($K_i$, $K_1$ and $k_3$) assessed region by region.

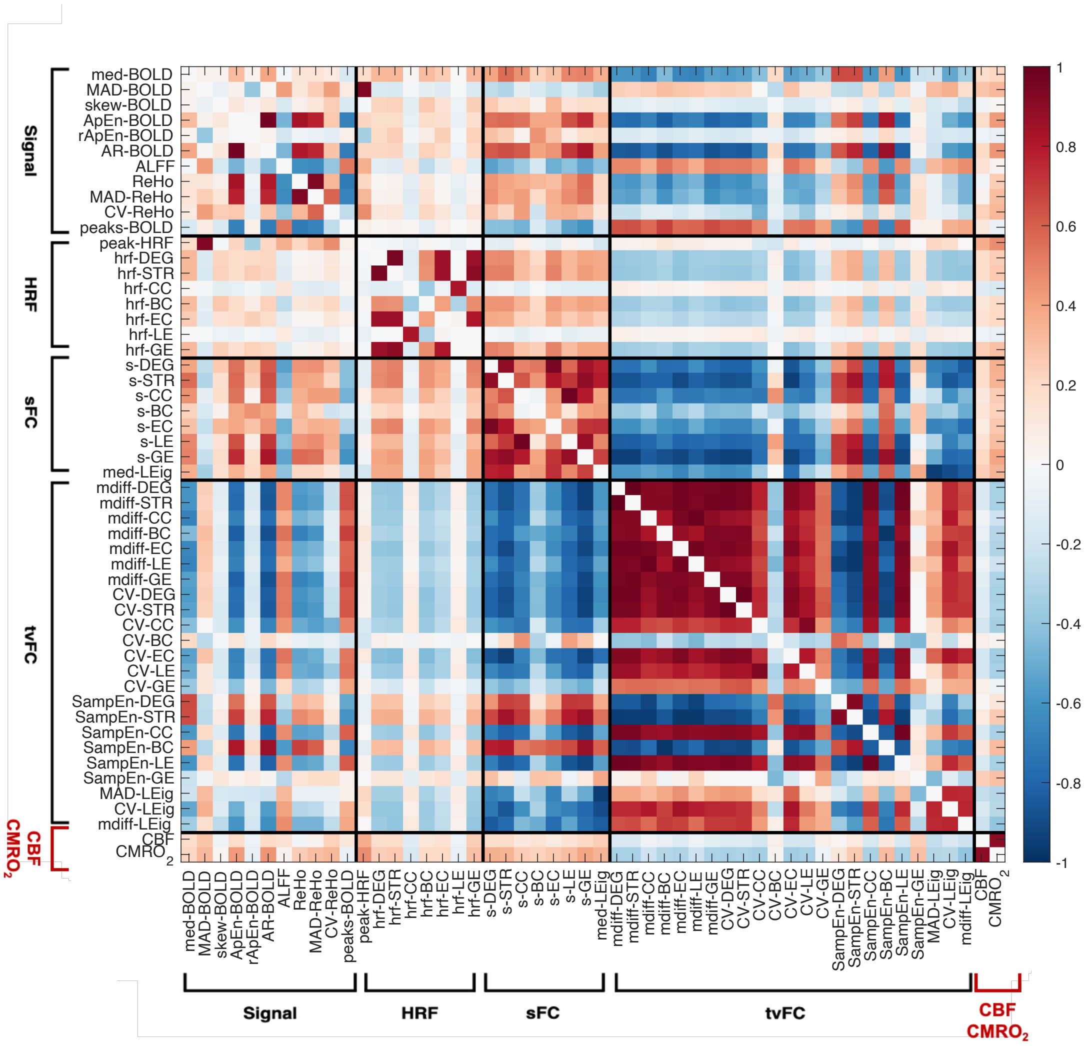


**Figure S6**: Pearson's correlations among group-average functional "predictor" variables, i.e., A) rs-fMRI features (divided into 4 a priori defined pools: 1) signal, 2) HRF, 3) sFC, 4) tvFC), B) CBF and $CMRO_2$.

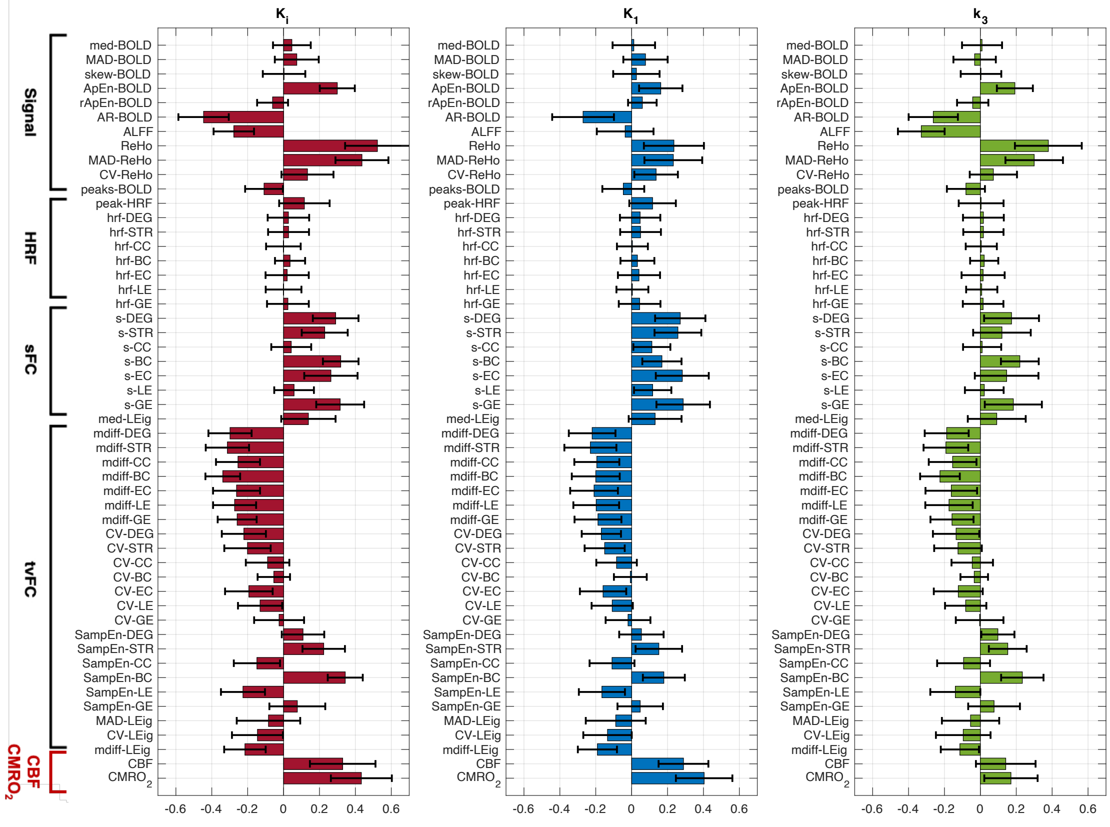


**Figure S7**: Pearson's correlations across brain regions of individual-level [$^{18}$F]FDG kinetic parameters ($K_i$, $K_1$ and $k_3$) with A) rs-fMRI features (divided into 4 a priori defined pools: 1) signal, 2) HRF, 3) sFC, 4) tvFC), B) CBF and $CMRO_2$. Mean and SD of r-to-z transformed correlations are reported.

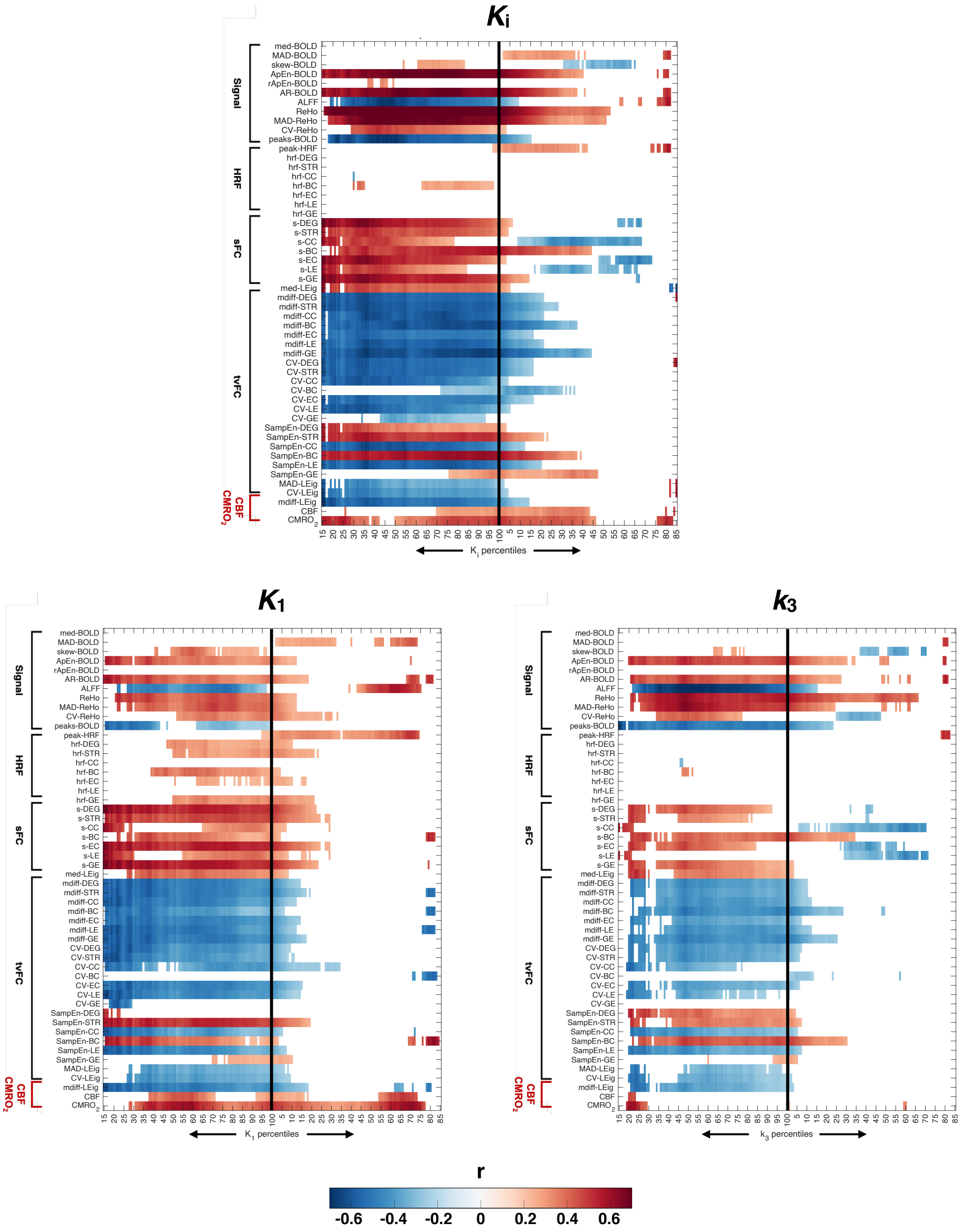


**Figure S8**: Pearson's correlations ($p < 0.05$, non-significant values in white) between [$^{18}$F]FDG kinetic parameters ($K_i$, $K_1$ and $k_3$) and A) all rs-fMRI features, B) CBF and $CMRO_2$ (*y axis*), evaluated across nodes selected by linearly increasing (*x axis – right side*) and decreasing (*x axis – left side*) percentiles of the $K_i$, $K_1$ and $k_3$ distributions.

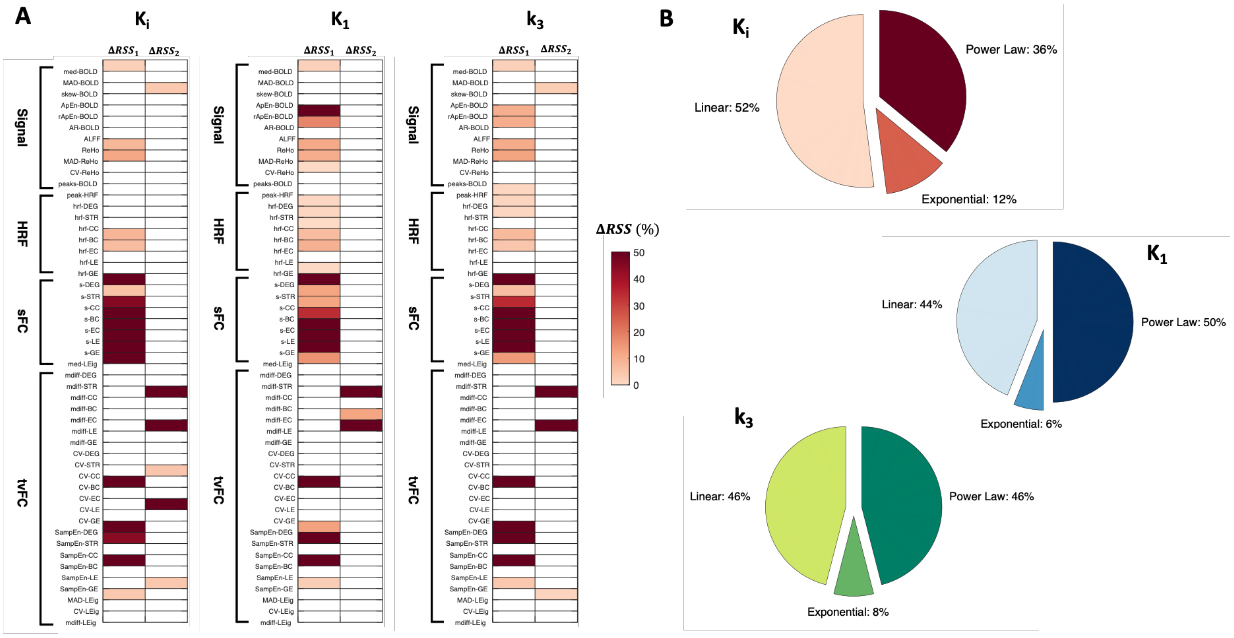


**Figure S9**: Assessment of nonlinearities in the bivariate spatial associations between [$^{18}$F]FDG parameters ($K_i$, $K_1$ and $k_3$) and rs-fMRI features: percentualized differences between linear and power model ($\Delta RSS_1$) and between linear and exponential model ($\Delta RSS_2$) for each rs-fMRI feature (**A**), and pie chart with the percentage of features (out of 50) whose association with $K_i$, $K_1$ and $k_3$ is best described by a linear, exponential, or power law model (**B**).

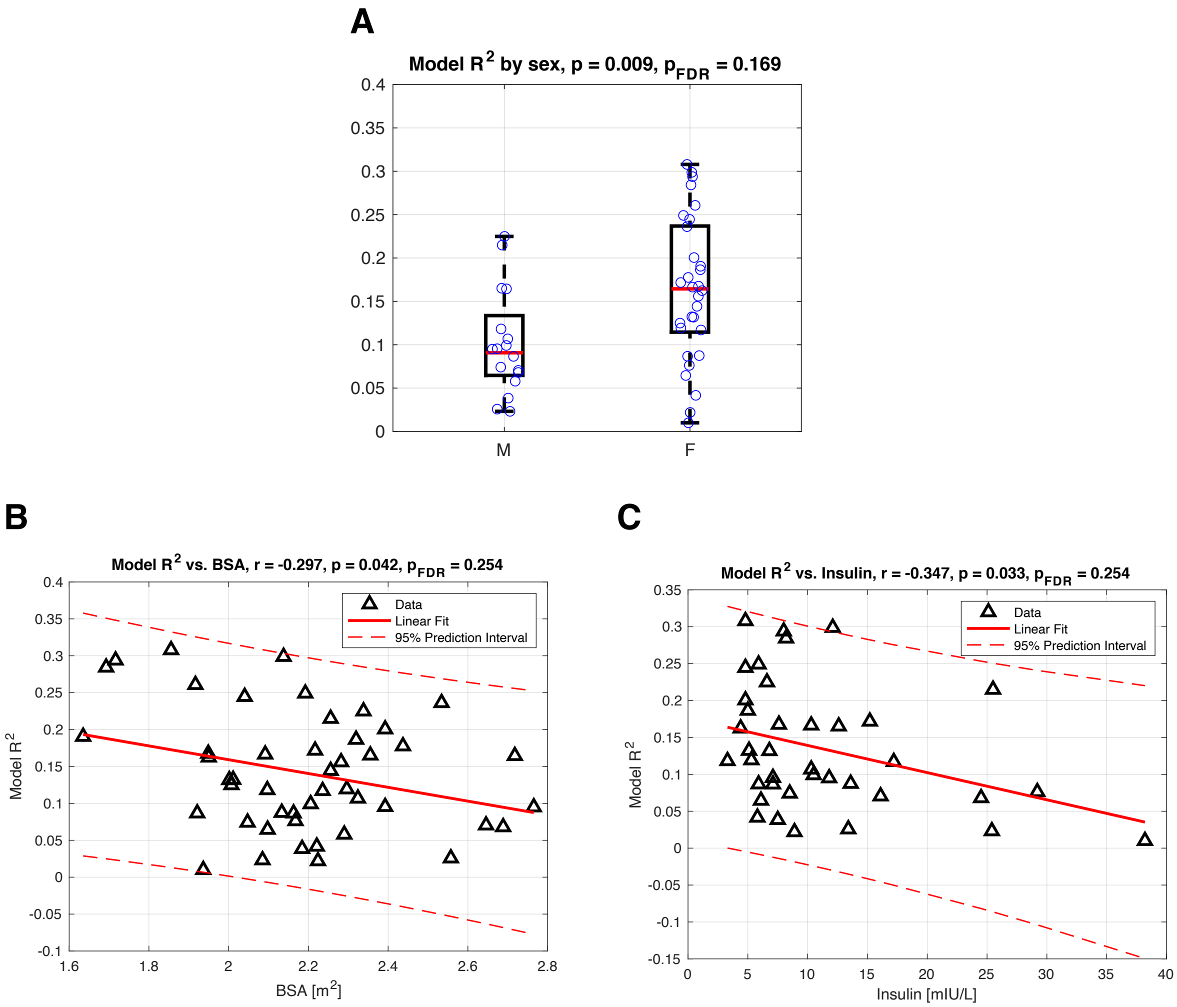


**Figure S10.** Multilevel modeling of $K_1$: individual variability vs. covariates in *Dataset 2*. Boxplots and scatter plots of the significant associations ($p<0.05$, uncorrected) between individual $R^2_i$ values of $K_1$ (adapted 9-parameter model) and participant covariates: sex (**A**), body-surface area – BSA [$m^2$] (**B**), insulin plasma levels [mIU/L] (**C**).

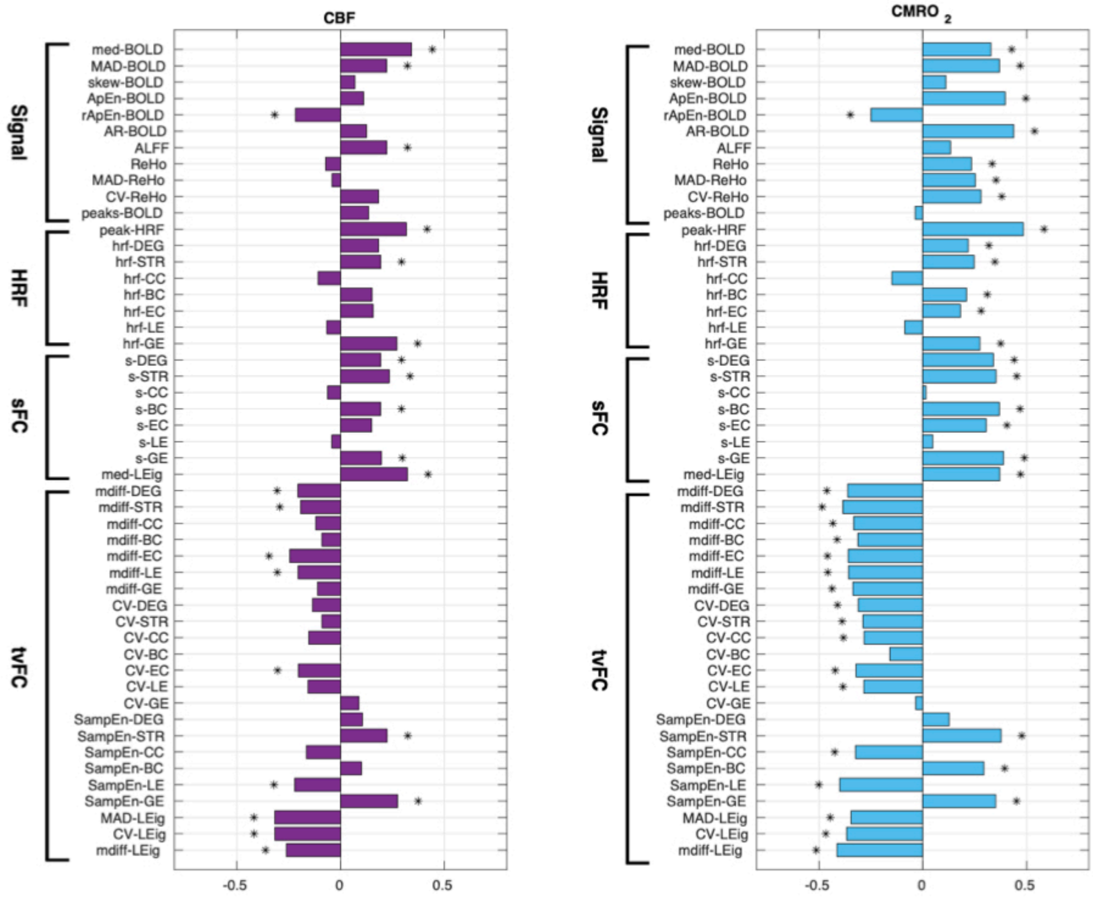


**Figure S11**: Pearson's correlations ($p < 0.05$, significant correlations highlighted with asterisk) across brain regions between group-average CBF and $CMRO_2$ vs. group-average rs-fMRI features.
